# Shifts in the Maternal Gut Microbiome Across Pregnancy are Associated with Energetic Status in Phayre’s Leaf Monkeys

**DOI:** 10.64898/2026.10.15.759161

**Authors:** Ting-Shuo Nien, Kiersten Grathwohl, Andreas Koenig, Carola Borries, Amy Lu, Elizabeth K. Mallott

## Abstract

**Background:** During pregnancy, mothers experience physiological changes that support fetal development, including shifts in energy utilization and increasing insulin resistance during late gestation. While these changes benefit the fetus, they can also impose energetic costs that affect the mother’s physical condition and metabolic phenotype. Previous studies suggest that the gut microbiome, a key regulator of female metabolism and energy harvest, may play a critical role in mediating these physiological challenges.

**Results:** In this study, metagenomic sequencing of the gut microbiome of 120 samples revealed dramatic shifts in gut bacterial composition and function associated with pregnancy and lactation in 14 wild Phayre’s leaf monkeys, *Trachypithecus phayrei crepusculus*, from two groups at the Phu Khieo Wildlife Sanctuary, Thailand. Alpha diversity, beta diversity, and taxonomic composition all differed between cycling and the 3^rd^ trimester. Furthermore, bacterial dissimilarity between these states was negatively correlated with postpartum physical condition; that is, mothers whose gut microbiota remained more stable during pregnancy recovered sooner after giving birth. Functional profiling revealed a depletion of pathways involved in purine salvage and glutamate degradation during the 3^rd^ trimester.

**Conclusion:** Overall, our study demonstrates that the gut microbiome undergoes substantial changes during pregnancy, and the stability of these microbial communities predicts postpartum maternal health and potentially influence lifetime reproductive output.

## Background

Human mothers experience numerous physiological changes during pregnancy to support fetal development. For instance, the maternal immune system undergoes significant changes to ensure the survival of the fetus [1]. In addition, maternal metabolism undergoes a significant transformation to meet the rising energetic demands during gestation and lactation [2]. These changes include increased fat storage and insulin sensitivity during early gestation to boost energy reserves, followed by enhanced fat utilization and higher insulin resistance during late gestation and lactation [3]. These patterns are also common in nonhuman primates. Research on wild female baboons (*Papio cynocephalus* and *P. anubis*) indicates that body fat increased throughout most of gestation but is depleted postpartum [4]. Similarly, both female common marmosets (*Callithrix jacchus*) and rhesus macaques (*Macaca mulatta*) gained body mass during pregnancy [5, 6]. These metabolic shifts, together with cardiovascular adaptations such as increased cardiac output and decreased vascular resistance, support the nutritional requirements of the growing fetus and neonate [7]; however, they can have negative consequences for the mother. For example, late-gestation insulin resistance is known to contribute to gestational diabetes mellitus (GDM) [8], and cardiovascular shifts are often linked to the development of preeclampsia [7].

These changes in maternal physiology during pregnancy and lactation are regulated by hormones [2, 9]. Estrogen and progesterone interact with the immune system during pregnancy by regulating Th cells and suppressing the adaptive immune response to promote immune tolerance [10]. Furthermore, these hormones contribute to the development of maternal insulin resistance, ensuring a steady nutrient supply for the developing fetus [10, 11]. Following birth, shifting hormone profiles become crucial for supporting milk production in early lactation [12]. Importantly, hormonal shifts during pregnancy and lactation may also impact the gut microbiota, which in turn may play a direct role in mediating the physiological changes during gestation [2]. For example, the gut microbiota can influence the immune response by regulating the balance between pro-inflammatory and anti-inflammatory cytokines and promoting the differentiation and expansion of Treg cells [13, 14]. In addition, the gut microbiota can impact metabolism by increasing the production of short-chain fatty acids, which stimulate the release of metabolic hormones, including glucagon-like peptide-1 (GLP-1) and peptide YY. These hormones affect insulin sensitivity and fat deposition, potentially shaping the metabolic environment of pregnancy and lactation [15]. The gut microbiota may also mitigate the energetic costs of both pregnancy and lactation by buffering mothers against nutrient shortfalls. In wild populations, the gut microbiota has been shown to compensate for seasonal variation in energy intake and dietary shifts [16, 17], as well as provide essential amino acids to support the metabolism [18]. Beyond hormonal shifts, dietary factors can also shape the gut microbiota [19], including changes in maternal energy intake due to seasonal foraging and the availability of specific phytoestrogen-rich foods like *Vitex,* which may affect endogenous progesterone activity [20].

Numerous studies have demonstrated changes in gut bacterial diversity throughout pregnancy and postpartum [21–24], as well as strong associations between gut microbial dynamics, female immunity, and metabolism. Yet research integrating microbial dynamics with hormones and measures of maternal energetics, health, and fitness remains limited in wild mammal populations. Furthermore, the majority of existing studies rely on 16S rRNA sequencing, which cannot provide information on changes in microbial function. Although several studies have employed metagenomic sequencing to examine pregnancy-associated changes in the gut microbiota, these efforts have been largely restricted to human populations [25–27], leaving a significant gap in our understanding of these processes in wild mammals.

To address these questions, we utilized a longitudinal paired dataset of metagenomic sequences and reproductive hormones from wild female Phayre’s leaf monkeys (*Trachypithecus phayrei crepusculus*), for which reproductive state was known. For a subset of these cases the metagenomic data could be paired with changes in maternal physical condition over time and the length of the subsequent interbirth interval (IBI), which were used as proxies for maternal energetic condition and recovery following pregnancy. Given the significant hormonal fluctuations across reproductive states and the established relationships between hormones and the gut microbiota [2], we predicted that fecal estrogen and/or progesterone concentrations would be associated with the composition and diversity of the gut microbiota. Similarly, we expected that shifts in maternal energy balance and the seasonal consumption of specific dietary resources like *Vitex* would be associated with gut microbial variation. In addition, using metagenomic data, we predicted an enrichment of functional pathways related to energy and nitrogen metabolism during both pregnancy and lactation to support the increased energetic and nutrient requirements. Finally, given the potential for the gut microbiota to buffer energetic and nutrient requirements, we hypothesized that variations in gut bacterial composition and function across reproductive states would be associated with maternal postpartum physical condition and IBI.

## Materials & Methods

### Study site and observational data collection

This study took place at Phu Khieo Wildlife Sanctuary (16°5’-35’N, 101°20’-55’E) in Chaiyaphum Province, Thailand as part of a long-term project on the behavioral ecology of wild Phayre’s leaf monkeys (directed by A.K. and C.B.). The study site is located 600-800m above sea level, and receives an average of 1186 mm of rain annually, with most rain falling during a wet season from April to October [28]. Reproduction is moderately seasonal in this population, with conceptions peaking from May to August, and most births falling between December and March [28]. The diet of Phayre’s leaf monkeys includes species from the genus *Vitex*, which produce phytoprogestins in both leaves and fruits that may influence the likelihood of conception [29]. Because the consumption of *Vitex* may affect endogenous progesterone activity, its availability represents a potential exogenous influence on maternal endocrine profiles and gut microbial dynamics [29] that we control for in the analysis (see below).

We collected fecal samples from two groups (PA and PB) between February 2005 and September 2006 as part of a study on reproductive physiology. Samples were collected from individually recognized adult females (PA: *N* = 4; PB: *N* = 10), stored in a cooler while in the field, and transferred to a –20°C freezer upon return to the field station (within 12 hours of sample collection). Samples were transported frozen to the United States, lyophilized, and sifted into fine powder at the Center for Conservation and Reproduction of Endangered Species in San Diego. We selected a subset of 120 fecal samples to determine gut microbiome composition. Females who had samples from at least two of three reproductive states (cycling, pregnant, or lactating) were included in the subset. Representative samples from each of the following reproductive periods were selected: cycling, 1^st^ trimester, 2^nd^ trimester, 3^rd^ trimester, lactation 1 (the first 3 months postpartum), and lactation 2 (4+5 months postpartum) (see **Table 1** for sample sizes). Lactation was partitioned to capture potential taxonomic variation in the gut microbiome occurring between the weeks immediately after birth and subsequent months [24, 30]. Samples from group PB (*N* = 101 from 10 females) were additionally analyzed for fecal hormone concentrations.

**Table 1.** Number of samples and individuals cycling, pregnant, and lactating.

|  | **Cycling** | **1<sup>st</sup>**<br>**trimester** | **2<sup>nd</sup>**<br>**trimester** | **3<sup>rd</sup>**<br>**trimester** | **Lactating 1** | **Lactating 2** |
| --- | --- | --- | --- | --- | --- | --- |
| **Number of**<br>**samples**<br>**(Number of**<br>**individuals)** | 22 (11) | 30 (11) | 34 (12) | 23 (8) | 5 (5) | 6 (6) |
| **Average**<br>**number of**<br>**samples per**<br>**individuals** | 2.0±0.6 | 2.7±1.0 | 2.8±0.4 | 2.9±0.8 | 1.0 | 1.0 |

All methods and protocols were approved by Stony Brook University IACUC (IDs: 20041120-20051120) and complied with the applicable laws of Thailand and the United States. Permission to conduct research in Thailand was granted by the National Research Council of Thailand, the Department of National Parks, Wildlife, and Plant Conservation, and Phu Khieo Wildlife Sanctuary.

For individuals from group PB, females were assigned as cycling, pregnant, or lactating based on the pattern of fecal estrogen (fE) and progesterone (fP) metabolites, which were analyzed as part of a previous study using radioimmunoassays [22, 29, 31]. These analyses allowed us to calculate the average length of gestation (205.3 days), the average cycle length (28.4 days), and the mean number of cycles (3.57 cycles) and days (101.4) to conception [31]. These data were then used to determine reproductive state for females in the second group, PA, for which fine-grained data on hormones were unavailable. For group PA, the beginning of lactation was defined by the observed date of birth; the beginning of gestation by the date of conception (calculated by subtracting the average gestation length from the observed date of birth); and the first day of cycling by subtracting the mean number of days until conception from the date of conception [31]. Data on the change in physical condition from pre- to post-partum were available from 6 females for which matching microbiome data were also available. Eileen Larney and C.B. assessed physical “condition monthly on a 7-point scale (1=meager; 7=fat) following previous methods [29, 32, 33]. This method has been successfully used in other studies of colobines to relate changes in body condition to seasonal variation in energy intake, energy expenditure, timing and probability of conception, and the rate of infant development[32, 34–37]. Interobserver reliability for the assessment of physical condition was nearly 100%. We calculated a z-score between physical condition at conception and at birth to determine changes over the course of pregnancy. Individuals with physical condition z-scores >0 were classified as having a better physical condition postpartum, while those with z-scores <0 were classified as having a worse physical condition postpartum. We supplemented the data on physical condition with data on the subsequent interbirth interval (IBI) if the current infant had survived the subsequent conception. It was available from 8 females. The IBI is defined as the time between two consecutive births. While this interval includes the subsequent gestation period, its overall length reflects the time required for a mother to recover from energy-draining lactation to return to a sufficient metabolic threshold for reconception [38, 39]. Thus, physical condition from conception to birth represents maternal energy storage during pregnancy and the IBI represents the duration of maternal energy recovery from lactation.

### DNA extraction and library preparation

We extracted DNA from lyophilized samples using a DNAeasy PowerLyzer Powersoil Kit (Qiagen) using established lab protocols [22]. The manufacturer’s protocol was modified by reducing the input sample weight to 0.1g, incubating samples at 65°C for 20 minutes after the addition of solution C1, and warming the solution C6 to 65°C prior to use. DNA libraries were prepared using the Illumina DNA Prep Kit and unique dual indexes (Illumina) following the manufacturers protocol for DNA inputs <100ng. We quantified libraries with a Qubit (Invitrogen) and verified fragment sizes on a TapeStation 4200 (Agilent). Libraries were equimolar pooled and sequenced on an Illumina NovaSeq platform (2×150bp) at the Genome Technology Access Center at Washington University in St. Louis. To account for potential background contamination, negative controls from the DNA extraction and library preparation steps were included during sequencing.

### Metagenomics sequencing and analysis

Our data analysis followed the pipeline described by Mallott et al. (2024) with minor modifications [40]. Briefly, metagenomic sequencing data were trimmed and filtered to remove low quality and potential host reads using KneadData (v0.12.0) with default settings [41]. The human reference genome hg37dec_v0.1 was used for host read removal. The filtered FASTQ files were taxonomically profiled using MetaPhlAn (v4.0.2) with the mpa_vJun23_CHOCOPhlAnSGB_202307 MetaPhlAn database to generate a species abundance table [41]. Functional profiling was performed using HUMAnN (v3.9) with the UniRef50_201901b protein database to generate a pathway abundance table [41]. Pathway abundance tables were normalized to counts per million and split into stratified and unstratified tables for further analysis.

### Microbial diversity and statistical analyses

Species richness, evenness, and Shannon diversity indices were calculated using the *vegan* package (v2.7-2) in R from the species abundance table [42]. Repeated measures correlation (*r_rm_*) was used to account for individual variation while testing the relationship between hormonal/environmental factors and gut microbiota diversity. This analysis was performed using the *rmcorr* package (v0.7.0), treating individual identity as the subject variable [43].

For beta diversity, Bray–Curtis and Jaccard distance matrices were computed using *vegan* (v2.7-2) from the species abundance table and the unstratified pathway abundance table [42]. Weighted UniFrac and unweighted UniFrac distance matrices were generated from the species abundance table using the *rbiom* package (v1.0.3) in R version 4.3.2 [44]. Beta diversity was visualized with nonlinear multidimensional scaling plots (NMDS). PERMANOVAs were used to evaluate the effects of reproductive state, individual identity, and group on bacterial and pathway composition using the Bray–Curtis, Jaccard, weighted UniFrac, and unweighted UniFrac distance matrices in *vegan* (v2.7-2) [42]. To control for the effects of environmental variation on the gut microbiome, we ran separate PERMANOVA models for each potential covariate: (i) expected monthly energy balance (high vs. low, based on published values of seasonal feeding time: Borries et al., 2024 [28]) and (ii) *Vitex* fruit availability. Maternal energy balance categories were applied to account for monthly variations in feeding time on high-quality foods and their expected effects on maternal energy balance. Months with expected positive energy balance were previously defined as the months following a three month or longer period of high young leaf and flower intake (April and May) and the months following a three month or longer period of high fruit intake (September–December) [28]. *Vitex* availability (April-September) serves as a proxy for potential phytoprogestin effects on endogenous hormones [29]. Posthoc pairwise multilevel comparisons among reproductive states for the PERMANOVA models were conducted using the *pairwiseAdonis* package (v0.4.1) [45].

Pearson correlation tests were used to examine associations between changes in physical condition and bacterial compositional change during pregnancy, using pairwise Bray–Curtis, Jaccard, weighted UniFrac, and unweighted UniFrac distances between the 3^rd^ trimester and cycling states. Similarly, Pearson correlation tests were used to examine the association between interbirth interval and bacterial compositional change during pregnancy, defined as the pairwise Bray–Curtis, Jaccard, weighted UniFrac, and unweighted UniFrac distances between the 3^rd^ trimester and cycling. Only individuals with available data on interbirth intervals or physical condition data were included in the correlation analyses. All analyses were performed in R version 4.5.1, unless otherwise specified [46].

### Identification of enriched taxa and metabolic pathways

We applied multiple complementary methods to identify taxa showing differential abundance across reproductive states, as these methods vary in their statistical assumptions, sensitivity, and ability to account for temporal structure in the data. Specifically, to identify taxa with significant variation across reproductive statereproductive states, we used a generalized linear mixed effects model with a negative binomial distribution. These models included individual identity as a random effect and were implemented using the *glmmTMB* (v1.1.13) and *car* (v3.1-3) packages in R [47, 48]. Tukey’s HSD was used for post-hoc comparisons using the *nlme* (v3.1-168) and *emmeans* (v2.0.2) packages [49, 50]. To identify potential key species distinguishing reproductive states, we focused on the 3^rd^ trimester and cycling, as these states showed significant differences in beta diversity based on pairwise multilevel comparisons from PERMANOVA models. We utilised a random forest algorithm using the *randomForest* package (v4.7-1.2) [51] to identify taxa that best discriminated between these periods. To identify potential taxa enriched or depleted in individuals with better or worse postpartum physical condition across reproductive states, we performed longitudinal analysis using *coda4microbiome* package (v0.2.4) [52]. Only individuals with available interbirth interval or physical condition data were included in the longitudinal analysis. To detect functional pathways enriched across reproductive states and between individuals with better or worse postpartum physical condition, we used the *limma* (v3.64.3) and *edgeR* (v4.6.3) packages in R, based on the unstratified pathway abundance table [53, 54]. Pathways with adjusted *p*-values < 0.05 and absolute log (fold change) >1 were considered significantly enriched or depleted.

### Network analysis

To compare microbial community connectivity across reproductive states and postpartum physical conditions, we performed network analysis using the *SpiecEasi* (v1.99.0) and *igraph* (v2.1.4) packages in R [55, 56]. The analysis pipeline followed Birt and Dennis., 2020 with minor modifications [57]. Briefly, taxa with very low prevalence (<1% of samples) were filtered from the abundance table to reduce network complexity. The filtered relative abundance table was then converted into a pseudo-count matrix. Networks were inferred from this matrix using the Meinshausen–Bühlmann neighborhood selection method. From the inferred networks, adjacency matrices were extracted to construct undirected, weighted graphs. Community structure (i.e., modules) was detected using the Louvain algorithm. Final node and edge tables, containing edge weights, module assignments, and interaction types (positive or negative), were generated for visualization and analysis in *Cytoscape* (v3.10.4) [58].

## Results

### Gut bacterial composition across reproductive states

The metagenomic sequencing yielded 11,516,958 ± 5,880,973 (range: 2,997,609 - 51,870,518) total reads and 3,478,121,320 ± 1,776,053,812 (range: 905,277,918 - 15,664,896,436) total bases per sample. Gut bacterial richness in the 3^rd^ trimester was significantly lower than in the 1^st^ trimester, lactation 2, and cycling (**Figure 1A, see Table S1 for complete pairwise comparisons**); a similar downward trend was observed compared to the 2^nd^ trimester. By contrast, evenness did not differ significantly among states (*F*(5, 101) = 0.53, *p* = 0.75; **Figure S1**). In addition, Shannon diversity, which gives greater weight to rare species, was significantly lower in the 3^rd^ trimester compared with the 1^st^ trimester, lactation 2, and cycling (**Figure 1B, Table S2**). Correlation analyses (*N* = 10 participants, *N* = 85 total observations) revealed negative correlations between fE levels and both species richness (*r_rm_* = −0.389, *p* < 0.001) and Shannon diversity (*r_rm_* = −0.429, *p* < 0.001) (**Figures 1C and 1D**). In addition, fP levels were negatively correlated with species richness (*r_rm_* = −0.256, *p* = 0.026) and Shannon diversity (*r_rm_* = −0.350, *p* < 0.010) (**Figure S2**)

**Figure 1.**
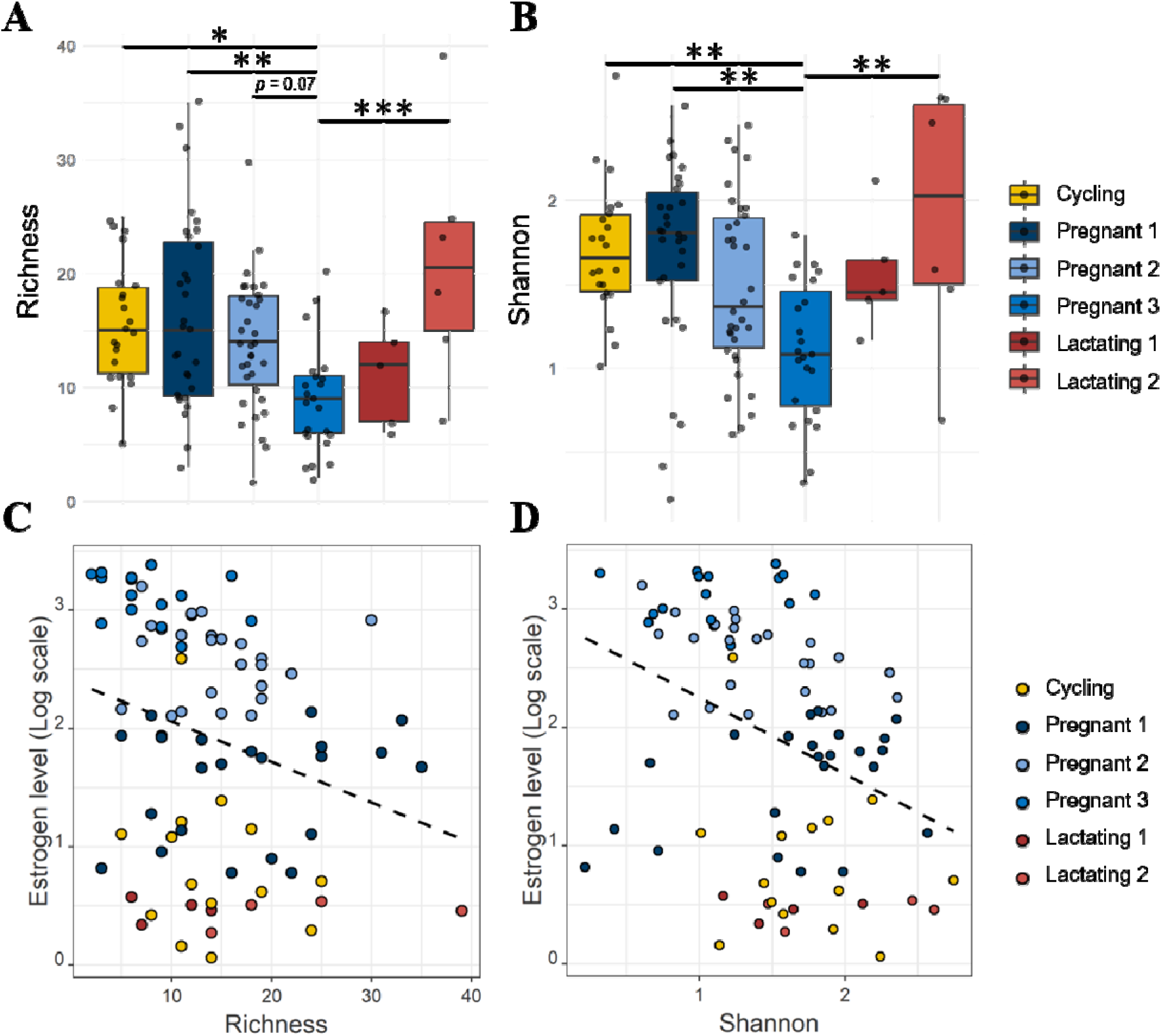
Species richness and Shannon diversity across reproductive states and their correlation with fE levels. Box plots showing (**A**) species richness and (**B**) Shannon diversity of gut microbiota in the cycling, 1^st^ (Pregnant 1), 2^nd^ (Pregnant 2), and 3^rd^ trimesters (Pregnant 3); and lactation (Lactating 1, i.e., month 1-3, Lactating 2, i.e., months 4+5). In each box plot, the center line represents the median, the box indicates the interquartile range (IQR), and the whiskers extend to 1.5 × IQR. Statistical significance levels are indicated as follows: \**p* < 0.05, \*\**p* < 0.01, \*\*\**p* < 0.001. Scatter plots showing correlations between fE levels and (**C**) species richness or (**D**) Shannon diversity. Dotted lines represent the linear regression fit. Repeated measures correlation coefficients (*r_rm_*) were *r_rm_*= −0.350 (*p* = 0.002) for species richness and *r_rm_* = −0.429 (*p* < 0.001) for Shannon diversity.

We found that reproductive state significantly predicted gut bacterial composition for all four distance matrices tested, with *R²* = 0.0645 (*F* = 1.75, *p* = 0.020) for weighted UniFrac (**Figure 2A**), *R²* = 0.0717 (*F* = 2.14, *p* = 0.001) for unweighted UniFrac (**Figure 2B**), *R²* = 0.0528 (*F* = 1.45, *p* = 0.010) for Bray–Curtis (**Figure S3A**), and *R²* = 0.0528 (*F* = 1.45, *p* = 0.027) for Jaccard distances (**Figure S3B**). Pairwise comparisons among reproductive states revealed that gut bacterial composition in the 3^rd^ trimester differed significantly from that in the 1^st^ trimester and cycling, based on weighted UniFrac (*R²* = 0.129, *F* = 6.38, *p* = 0.015)., unweighted UniFrac (*R²* = 0.0792, *F* = 3.70, *p* = 0.015). and Jaccard distances (*R²* = 0.0651, *F* = 2.99, *p* = 0.015). Moreover, differences were observed between the 3^rd^ and 2^nd^ trimesters based on unweighted UniFrac (*R²* = 0.0670, *F* = 3.93, *p* = 0.015) and Jaccard distances (*R²* = 0.0469, *F* = 2.71, *p* = 0.030). Correlation analyses showed that NMDS1 from the weighted UniFrac exhibited no correlation with fE (*r_rm_* = 0.175, *p* = 0.10), but a significant positive correlation with fP (*r_rm_*= 0.251, *p* = 0.020) (**Figures 2C and S4A**). By contrast, NMDS1 from the unweighted UniFrac was weakly negatively correlated with fE (*r_rm_* = −0.262, p = 0.013), with no significant correlation observed with fP (**Figures 2D and S4B**).

**Figure 2.**
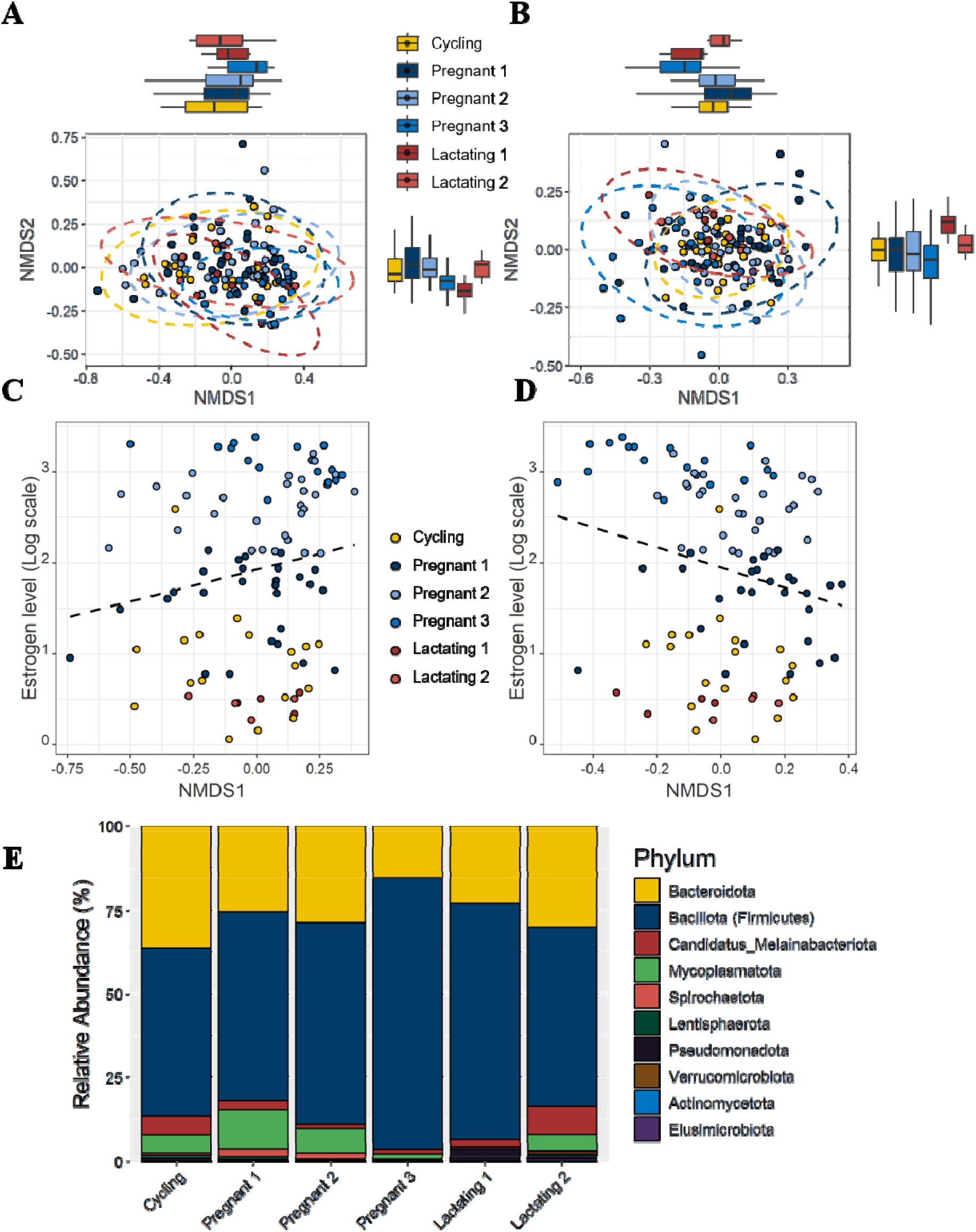
Shifts in gut microbial community structure and composition across reproductive states. Nonlinear multidimensional scaling (NMDS) plots based on (**A**) weighted UniFrac and (**B**) unweighted UniFrac of gut microbiota across reproductive states. Box plots along the top and right sides represent the distribution of NMDS1 and NMDS2 values for each reproductive state. In each box plot, the center line represents the median, the box indicates the interquartile range (IQR), and the whiskers extend to 1.5 × IQR. For the weighted UniFrac NMDS plot, stress = 0.163, *R²* for reproductive states = 0.065 (*p* = 0.020). For the unweighted UniFrac NMDS plot, stress = 0.205, *R²* for reproductive states = 0.072 (*p* = 0.001). Scatter plots showing correlations between fE levels and NMDS1 values derived from (**C**) weighted UniFrac and (**D**) unweighted UniFrac. Dotted lines represent the linear regression fit. Repeated measures correlation coefficients (*r_rm_*) were *r_rm_ =* 0.175 (*p* = 0.10, ns) for the weighted UniFrac and *r_rm_* = -0.262 (*p* = 0.013) for the unweighted UniFrac. (**E**) Stacked bar chart of the average relative abundance of microbial phyla across reproductive states.

### The effect of environmental factors on gut bacterial composition

We found that expected maternal energy balance was a significant predictor of bacterial composition based on unweighted UniFrac (*R²* = 0.0123, *F* = 1.85, *p* = 0.034), but not weighted UniFrac. Reproductive state remained a significant predictor of community composition across both unweighted UniFrac (*R²* = 0.0742, *F* = 2.23, *p* = 0.001) and weighted UniFrac (*R²* = 0.0629, *F* = 1.70, *p* = 0.024) models, even after accounting for individual identity, group ID, and variation in expected energy balance (**Figures 2A and 2B**).

In order to determine whether reproductive state remained a robust predictor of microbial dynamics even when accounting for the consumption of a plant containing phytohormones, which could affect maternal endocrine profile [29], we used a PERMANOVA model that included both *Vitex* availability and reproductive state, along with individual ID and group ID as predictors. We found that *Vitex* availability was a significant predictor of bacterial composition based on weighted UniFrac (*R²* = 0.0226, *F* = 3.12, *p* = 0.013) and unweighted UniFrac (*R²* = 0.0410, *F* = 6.42, *p* = 0.001). Importantly, reproductive state remained a significant predictor of community composition based on weighted UniFrac (*R²* = 0.0616, *F* = 1.70, *p* = 0.018) and unweighted UniFrac (*R²* = 0.0584, *F* = 1.83*, p* = 0.001), even after accounting for the effect of *Vitex* consumption. Comparisons of hormone levels between samples collected during the *Vitex* season and non-*Vitex* season showed that fE levels were significantly lower during the *Vitex* season **(Figure S5A**), whereas fP concentrations were significantly higher **(Figure S5B**).

### Variation in phyla and taxa across reproductive states

After controlling for multiple comparisons, only the abundance of the phylum Spirochaetota differed significantly across reproductive states (between the 3^rd^ and 2^nd^ trimesters (*z* = -3.49, *p* = 0.006), between the 3^rd^ trimester and lactation 2 (*z* = -2.97, *p* = 0.033), and between the 2^nd^ trimester and cycling (*z* = 3.14, *p* = 0.019) (Tukey’s HSD; **Figure 2E**). At the species level, the GLMM with a negative binomial distribution did not identify any species with statistically significant differences across reproductive states (**Table S3**). To further identify taxa that were important in distinguishing the 3^rd^ trimester from cycling, we applied a Random Forest algorithm to classify samples from these two states (**Figure S6A**). The model demonstrated a high degree of discriminatory power with an Area Under the Curve (AUC) of 0.911. Specifically, the model classified samples from cycling with an error rate of 27% and samples from the 3^rd^ trimester with an error rate of 9% (**Figure S6B**). The six taxa with the greatest predicted value, as measured by the highest mean decrease in accuracy (MDA) were selected for comparison of their relative abundances across reproductive states (**Figure S7)**. Three taxa belonged to the phylum Bacteroidota (GGB1672_SGB2288, *Phocaeicola_dorei*, GGB13041_SGB112132), two to Bacillota (GGB3185_SGB4208, GGB9717_SGB15270), and one to Candidatus Melainabacteriota (GGB86634_SGB115439). Notably, GGB9717_SGB15270 was significantly more abundant in the 3^rd^ trimester compared to both the 2^nd^ trimester and cycling (**Figure S7**). The remaining five species did not exhibit statistically significant differences across states, despite their identified importance in the Random Forest classification model (**Figure S6**).

To evaluate whether microbial interactions differed between the 3^rd^ trimester and cycling, we performed a network analysis to assess network complexity (**Figures S8A and S8B**). The microbial network during cycling contained more nodes (39 vs. 19) and edges (31 vs. 11) than the network in the 3^rd^ trimester. In addition, the cycling network exhibited higher density (0.013 vs. 0.011) and a longer average path length (1.98 vs. 1.01) compared with the 3^rd^ trimester (**Figure S8C**). Taken together, these results indicate that gut microbial interactions were more complex during cycling than during the 3^rd^ trimester.

### Functional profiling of gut microbiome across reproductive states

We revealed significant differences in microbial functional pathways among reproductive states, with *R²* = 0.0967 (*F* = 2.98, *p* = 0.003) for Bray–Curtis distances (**Figure 3A**) and *R²* = 0.058 (*F* = 1.52, *p* = 0.033) for Jaccard distances (**Figure S9A**). When comparing differentially abundant pathways between the 3^rd^ trimester and cycling, 28 pathways were significantly enriched and 10 pathways were significantly depleted in the 3^rd^ trimester relative to cycling, with log_2_|fold change| > 1 and adjusted *p*-value < 0.05 (**Figure 3B**). The heatmaps further demonstrate the distinct patterns of pathway abundance between the 3^rd^ trimester and cycling (**Figures 3C and S9B**).

**Figure 3.**
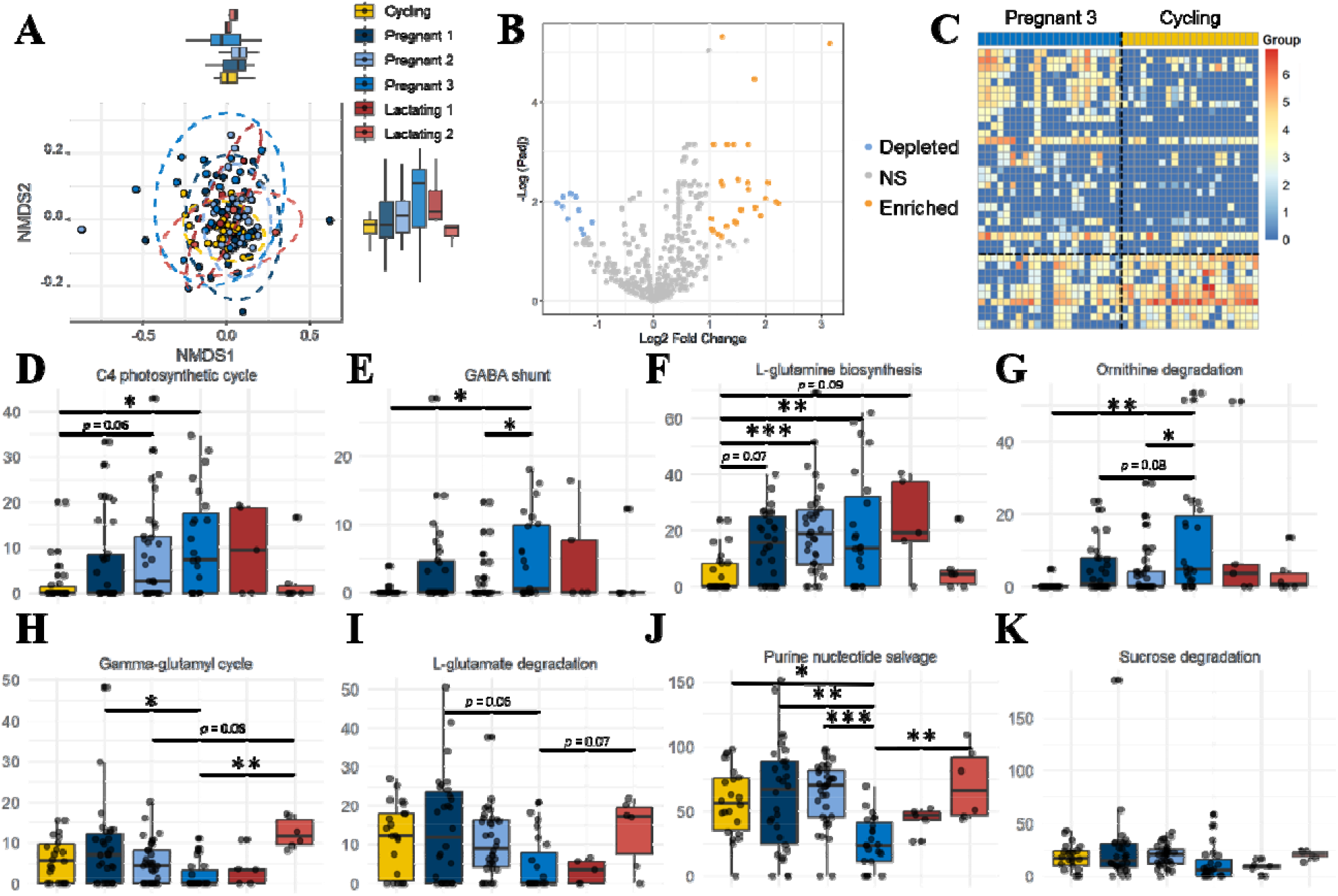
Functional profiling of gut microbiota across reproductive states. (**A**) Nonlinear multidimensional scaling (NMDS) plot based on Bray–Curtis distances of gut microbiota pathway abundances across reproductive states. Box plots along the top and right sides show the distribution of NMDS1 and NMDS2 values for each reproductive state. Stress = 0.126; *R²* for reproductive states = 0.097 (*p* = 0.003). (**B**) Volcano plot of functional pathways comparing the 3^rd^ trimester and cycling. Orange and blue indicate enriched and depleted pathways (adjusted *p* <0.05); NS, non-significant. (**C**) Heatmap of pathways enriched or depleted in the 3^rd^ trimester compared with cycling. The color scale represents the logarithm of pathway read counts, with red indicating enrichment and blue indicating depletion in the 3^rd^ trimester. (**D-K**) Box plots showing the relative abundance of selected pathways across reproductive states. In each box plot, the center line represents the median, the box indicates th interquartile range (IQR), and the whiskers extend to 1.5 × IQR. Statistical significance levels are indicated as follows: \**p* < 0.05, \*\**p* < 0.01, *** *p* < 0.001.

Several pathways showing differential abundance were compared across all reproductive states (**Figures 3D-K, Table S4 - S11**). Specifically, pathways involved in carbohydrate and amino acid metabolism—including C4 photosynthetic cycle, GABA shunt, L-glutamine biosynthesis, and ornithine degradation—were significantly enriched in the 3^rd^ trimester compared with cycling (**Figures 3D-G, Table S4 – S7**). By contrast, pathways involved in amino acid transport and nucleotide recycling, such as γ-glutamyl cycle and purine nucleotide salvage were significantly depleted in the 3^rd^ trimester compared with the 1^st^ trimester or cycling (both adjusted *p* < 0.05) (**Figures 3H and 3J, Table S8 and S9**). While L-glutamate degradation showed a non-significant trend toward depletion in the 3^rd^ trimester relative to cycling (adjusted *p* = 0.07) and lactating 2 (adjusted *p* = 0.07) (**Figure 3I, Table S10**), sucrose degradation, a primary carbohydrate metabolism pathway, exhibited no statistically significant differences across the reproductive cycle after correction for multiple comparisons (**Figure 3K, Table S11**).

### Association of gut microbiota with proxies of energetic status and recovery

Six individuals with metagenomic sequencing data from both the 3^rd^ trimester and cycling were included in a correlation analysis to examine the relationship between changes in physical condition and the gut microbiota across reproductive states. Among these individuals, three showed improved condition postpartum (Physical condition score > 0), one showed worsened condition (Physical condition score < 0), and two exhibited no change (Physical condition score = 0). We found a moderate negative correlation between changes in physical condition and the degree of bacterial compositional dissimilarity between the 3^rd^ trimester and cycling. Pearson’s correlation coefficients were *r* = -0.493 (*p* = 0.038) for weighted UniFrac (**Figure 4A**), *r* = −0.391 (*p* = 0.11) for unweighted UniFrac (**Figure 4B**), *r* = −0.521 (*p* = 0.027) for Bray–Curtis (**Figure 4C**), and *r* = −0.491 (*p* = 0.039) for Jaccard distances (**Figure 4D**). In other words, individuals whose gut microbiota differed more between the 3^rd^ trimester and cycling tended to have worsened physical condition from conception to the end of pregnancy.

**Figure 4.**
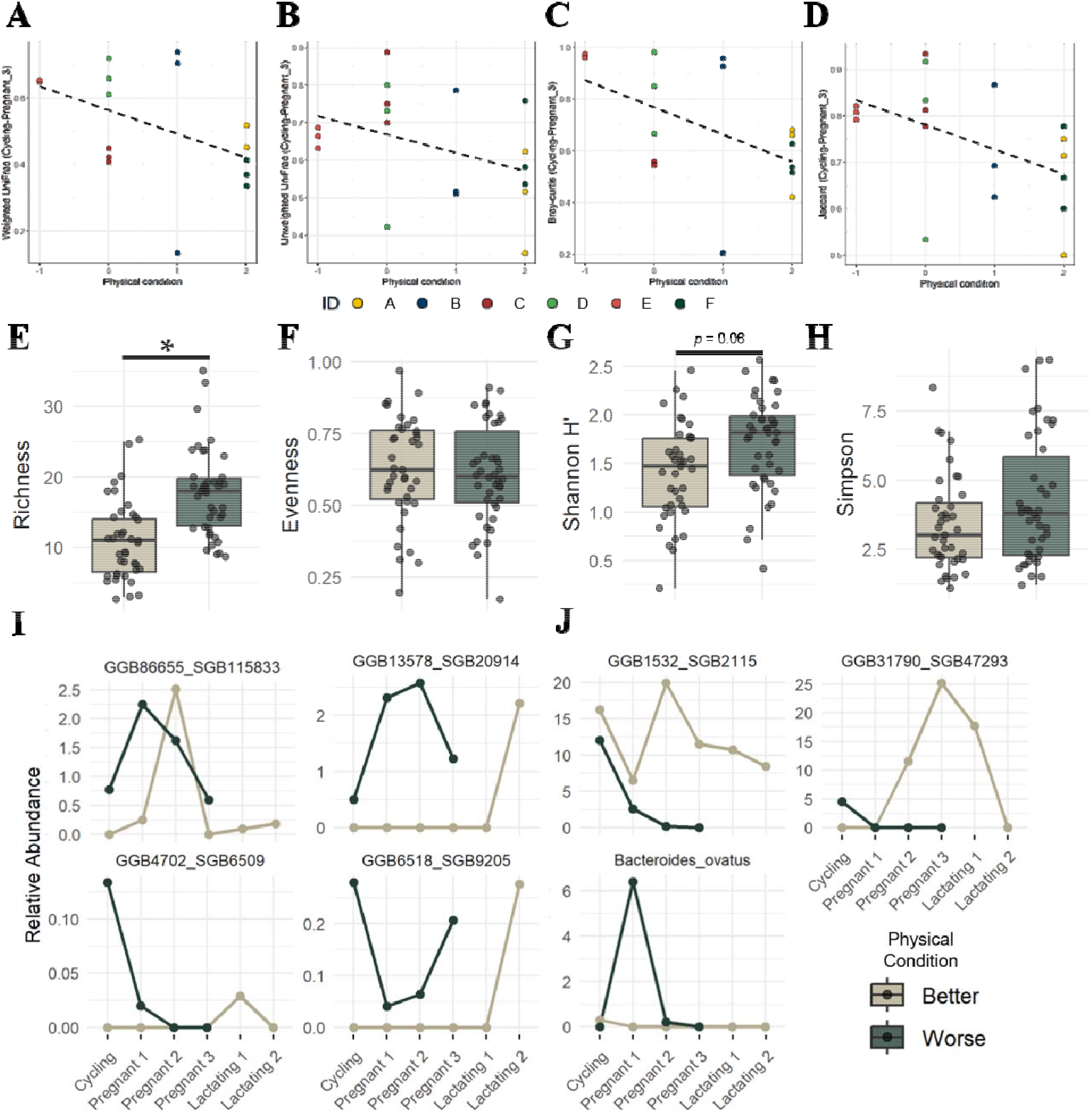
Association between gut microbiota composition and postpartum physical condition. (**A-D**) Scatter plots showing correlations between physical condition and species dissimilarity between the cycling and 3^rd^ trimester. Species dissimilarity was calculated using (**A**) weighted UniFrac, (**B**) unweighted UniFrac, (**C**) Bray–Curtis, and (**D**) Jaccard distances. Dotted lines represent the linear regression fit. Pearson’s correlation coefficients were *r* = -0.493 (*p* = 0.038) for weighted UniFrac, *r* = −0.391 (*p* = 0.11, ns) for unweighted UniFrac, *r* = −0.521 (*p* = 0.027) for Bray–Curtis, and *r* = −0.491 (*p* = 0.039) for Jaccard distances. Each ID represents an individual monkey. (**E-H**) Box plots showing (**E**) species richness, (**F**) evenness, (**G**) Shannon diversity, and (**H**) Simpson diversity between individuals with better and worse physical condition. Statistical significance levels are indicated as follows: \**p* < 0.05. Longitudinal analysis across reproductive states showing taxa (**I**) depleted or (**J**) enriched in individuals with better versus worse physical condition.

Furthermore, species richness was significantly lower in individuals with improved physical condition compared to those with worsened condition (*t*(6) = 3.06; *p* = 0.02; **Figure 4E**). By contrast, no significant differences were observed in Shannon diversity (*t*(6) = 2.35; *p* = 0.06), species evenness (*t*(6) = -0.34; *p* = 0.74), or Simpson diversity (*t*(6) = 1.82; *p* = 0.12) between the two groups (**Figures 4F-4H**). Longitudinal analysis revealed several taxa whose relative abundances significantly differed based on postpartum physical condition (**Figure 4I**). Specifically, GGB86655_SGB115833 (Spirochaetota, Brachyspiraceae), GGB13578_SGB20914, GGB4702_SGB6509 (Bacillota), and GGB6518_SGB9205 (Lentisphaerota) exhibited a general pattern of depletion in individuals with better condition, particularly during pregnancy. Conversely, GGB1532_SGB2115 (Bacteroidota), GGB31790_SGB47293 (Bacillota), and *Bacteroides ovatus* (Bacteroidota) maintained higher overall abundances in individuals with better condition (**Figure 4J**). Network analyses combining data across all reproductive states and comparing individuals with better versus worse condition showed that gut microbial interactions were more complex in individuals whose physical condition worsened between cycling and the last trimester (**Figures S10A and S10B**). Specifically, their microbial network contained more nodes (82 vs. 27) and edges (75 vs. 17), exhibited higher density (0.014 vs. 0.010) and a longer average path length (5.34 vs. 1.42), and showed a higher clustering coefficient (0.129 vs. 0) compared with individuals in better condition (**Figure S10C**). These results suggest that worsened postpartum physical condition is associated with a more complex gut microbial network.

Among the eight individuals included in the analysis, the mean IBI was 23.6 ± 4.5 (range: 15.1-28.0) months. To determine if the magnitude of the microbiome shift during pregnancy predicts future reproductive timing, we assessed four individuals with metagenomic sequencing data from both cycling and the 3^rd^ trimester. There was no significant correlation between IBI length and the degree of bacterial compositional dissimilarity between these states (**Figure S11**).

### Functional profiling of gut microbiome from individuals with better or worse postpartum physical condition

Twenty-three pathways were significantly enriched, and 24 pathways were significantly depleted in individuals with better physical condition relative to those with worsened condition (log_2_|fold change| > 1, adjusted *p* < 0.05; **Figure 5A**). Heatmaps further highlight distinct patterns of pathway abundance between the two groups (**Figures 5B and S12**). Several pathways related to amino acid and carbohydrate metabolism showing differential abundance were further compared across the two groups (**Figures 5C–5J**). Pathways involved in ornithine degradation, the C4 photosynthetic cycle, geranylgeranyl diphosphate biosynthesis, and the TCA cycle were significantly enriched in individuals with better physical condition (all adjusted *p* < 0.05) (**Figures 5C–5F**). By contrast, pathways involved in L-glutamate degradation, purine nucleotide salvage, and sucrose degradation were significantly depleted in the improved-condition group compared to the worsened-condition group (all adjusted *p* < 0.05) (**Figures 5G–5J**).

**Figure 5.**
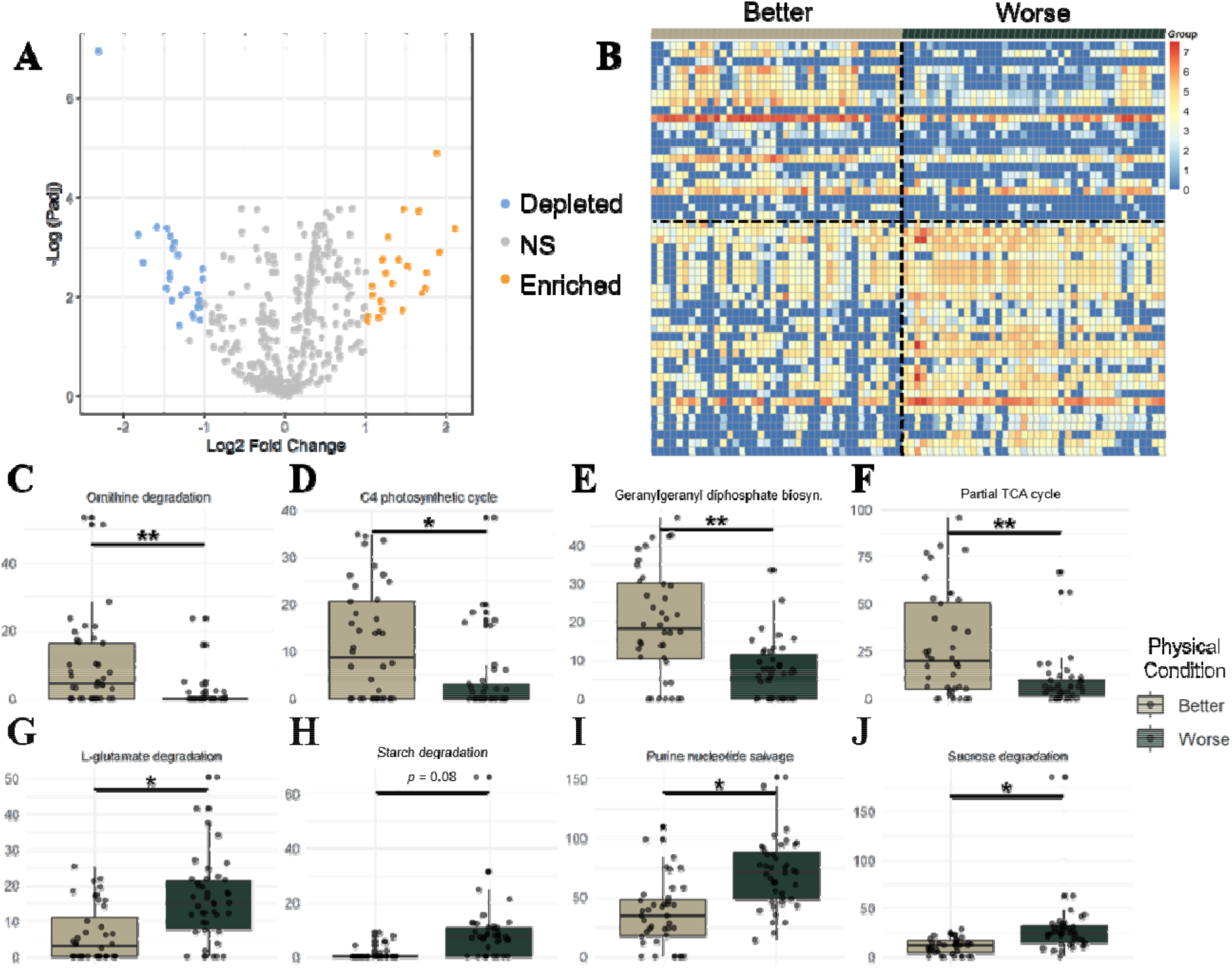
Functional profiling of gut microbiota between individuals with better and worse postpartum physical condition. (**A**) Volcano plot of functional pathways comparing individuals with better versu worse physical condition. Orange and blue indicate enriched and depleted pathways (adjusted *p* < 0.05); NS, non-significant. (**B**) Heatmap of pathways enriched or depleted in individuals with better compared to worse physical condition. The color scale represents the logarithm of pathway read counts, with red indicating enrichment and blue indicating depletion in the 3^rd^ trimester. (**C-J**) Box plots showing th relative abundance of selected pathways across different levels of physical condition. In each box plot, the center line represents the median, the box indicates the interquartile range (IQR), and the whiskers extend to 1.5 × IQR. Statistical significance levels are indicated as follows: \**p* < 0.05, \*\**p* < 0.01.

## Discussion

We found that the composition and function of the gut microbiome differ significantly across reproductive states in female Phayre’s leaf monkeys. It suggests that larger changes in the gut microbiome over the course of pregnancy are negatively associated with health- and fitness-relevant traits. The 3^rd^ trimester showed notably lower alpha diversity compared with cycling. Beta diversity analyses also consistently revealed significant differences in gut bacterial composition between the 3^rd^ trimester and cycling, regardless of the metric of analysis. Consistent with previous studies, differences in alpha and beta diversity were associated with fP and fE concentrations [22]. In addition, we identified specific taxonomic and functional signatures of late gestation, including the depletion of the Spirochaetota and significant enrichment in pathways related to carbohydrate and amino acid metabolism.

Our findings demonstrate that the Phayre’s leaf monkey gut microbiome undergoes significant structural reorganization during pregnancy, specifically characterized by a decrease in alpha diversity and a distinct shift in beta diversity during the third trimester. These results are consistent with our previous 16S rRNA study in this same population, which similarly reported a decrease in species richness and Shannon diversity during pregnancy compared with cycling individuals, along with shifts in gut microbial composition [22]. Furthermore, our results mirror patterns observed in other mammals, although the direction and magnitude of these shifts vary across species [59–68]. Multiple studies in mice (*Mus musculus*) using 16S rRNA sequencing have reported differences in gut bacterial composition across reproductive states or between pregnant and non-pregnant individuals, although at least one study observed an increase in the Shannon diversity during pregnancy [62–64]. Studies in humans [23, 24, 65, 66] and in several species of nonhuman primates using 16S rRNA sequencing have reported mixed results [19, 21, 22, 67–69]. For instance, a study of Tibetan macaques (*M. thibetana*) reported significant changes in gut bacterial composition across cycling, pregnancy, and lactation, but no differences in alpha diversity [67]. Research on yellow (*Papio cynocephalus*) and Anubis baboons (*P. anubis*) found an increase in ASV richness during pregnancy and significant differences in beta diversity throughout the reproductive cycle [68]. A study of geladas (*Theropithecus gelada*) found no significant differences in either alpha or beta diversity across reproductive states but nonetheless revealed that the relative abundances of *Verrucomicrobiota* and *Epsilonbacteraeota* were significantly higher during pregnancy than during cycling or lactation [19]. Finally, in white-faced capuchins (*Cebus imitator*), one study identified associations between reproductive state and gut bacterial composition [21], while another study reported that reproductive state was not a significant predictor of either Chao1 richness or gut microbial community dissimilarity [69]. The mixed results across nonhuman primate studies suggest that reproductive shifts in gut microbial diversity are species- or population-dependent, possibly driven by differences in maternal investment strategies and/or the seasonality of environments and food availability or lack thereof. For instance, to meet the energetic demands of gestation and lactation, primate females may adjust their behavior, increase energy intake and/or rely on stored energy reserves [70]. Higher levels of microbial plasticity may help compensate for energetic shortfalls during reproduction, both for primate species that increase their energy intake during reproduction as well as those who may rely on microbial changes to increase the efficiency of energy extraction from the diet during pregnancy and lactation. Moreover, these mixed results may also reflect differences in sampling methods or data analysis approaches among studies. Further research should consider how the timing of conception and birth aligns with seasonal fluctuations in food availability, as well as include additional primate species, to better understand the causes of this variation.

Consistent with previous studies, including one using 16S sequencing from the same study population [22], our results suggest that hormones such as estrogen and progesterone may play major roles in shaping the gut microbiome during gestation. Specifically, we found that maternal hormonal dynamics were strong predictors of gut microbial shifts, with both fE and fP concentrations negatively correlated with alpha diversity. Furthermore, fP concentrations significantly predicted differences in community composition based on weighted UniFrac. The relationship between reproductive hormones and the gut microbiome is further supported by studies in other mammals, although the direction of these effects appears to be species-specific. For instance, in contrast to the negative correlations observed here, fE and fP concentrations in baboons were positively correlated with ASV richness and significantly associated with variation in beta diversity [68]. In the eastern black rhino (*Diceros bicornis michaeli*), several gut microbial genera correlated significantly with fP metabolite concentrations [71]. Furthermore, experimental studies have shown that progesterone supplementation in mice, or incubation of fecal samples with progesterone, increased the relative abundance of *Bifidobacterium* [72]. Collectively, although the direction of hormone–diversity associations varied across studies and systems, these findings support a close relationship between hormonal dynamics and gut microbial diversity during pregnancy.

We also observed significant differences in microbial function across reproductive states. Specifically, pathways involved in the GABA shunt, ornithine degradation, and L-glutamine biosynthesis were significantly enriched in the 3^rd^ trimester compared with cycling. The GABA shunt pathway is not only crucial for the synthesis of γ-aminobutyric acid (GABA) from glutamate, but it also contributes to cellular energy metabolism by supplying NADH and succinate to the electron transport chain [73]. Gut microbiota, such as Bacteroidota species, may contribute to the intestinal GABA pool, potentially linking microbial metabolism to the gut–brain axis and host mood and behavioral regulation [74, 75]. Ornithine degradation provides an alternative route to GABA production via polyamine metabolism. In this pathway, glutamate also contributes to GABA synthesis indirectly through reactions involving glutathione metabolism and ornithine-derived polyamines [73]. In addition, L-glutamine biosynthesis is tightly linked to glutamate metabolism, as glutamate serves as the immediate precursor for glutamine through amidation catalyzed by glutamine synthetase [76]. Because these pathways converge on glutamate, our results suggest that glutamate metabolism becomes increasingly important during the 3^rd^ trimester, potentially supporting enhanced GABA and glutamine production. Glutamate serves as a central hub for amino acid metabolism, and glutamate-derived glutamine is critical for meeting the intense metabolic demands of the developing fetus and placenta [77]. Supporting this hypothesis, our study also revealed depletion in the γ-glutamyl cycle in the 3^rd^ trimester compared with 1^st^ trimester. This pathway is involved in the synthesis and degradation of glutathione, for which glutamate is a primary precursor [78], potentially allowing more glutamate to be available for glutamine production. In addition to glutamate-related metabolic pathways, our results also showed that pathway involved in purine nucleotide salvage was depleted during the 3^rd^ trimester compared with cycling. This depletion may reflect the increased maternal demand for nitrogen during late pregnancy, leaving fewer of these essential resources for microbes to degrade. However, it is important to note that these interpretations are based on metagenomic analysis, and we cannot directly confirm whether these pathways were actively expressed. Moreover, the observed enrichment or depletion of pathways may reflect shifts in microbial taxonomic composition rather than true changes in metabolic activity. Further studies integrating functional assays, metatranscriptomic analyses, or metabolomic data are needed to clarify how pregnancy-associated changes in gut microbiota composition both change microbial function and differentially influence female metabolic function across reproductive states.

This study also assessed the association of gut microbiota with measures of maternal physical condition. Our results suggest that individuals whose gut microbiota changed the most during pregnancy tended to exhibit worsened postpartum physical condition. Consistent with this interpretation, several studies have reported associations between disrupted gut microbial composition and adverse maternal health outcomes in humans, including GDM, early-onset preeclampsia and postpartum depressive disorder [79–82]. Furthermore, we evaluated maternal energetic recovery following birth, proxied via interbirth interval (IBI). We found no significant association between IBI and microbiota shifts during the transitions from cycling to the 3^rd^ trimester. While this implies that transient gestational microbial shifts may not directly predict subsequent reproductive timing, IBI is a rough proxy for maternal energy recovery. Given this limitation and our small sample size, the use of a more reliable physiological marker for energy status, such as urinary C-peptide [83], and larger datasets are required to fully evaluate these relationships. In addition, we identified several taxa that were enriched or depleted in individuals with better postpartum health (Taxa from Bacillota (GGB13578_SGB20914, GGB4702_SGB6509), Lentisphaerota (GGB6518_SGB9205), and Spirochaetota (GGB86655_SGB115833) were depleted in individuals with better physical condition, whereas taxa from Bacteroidota (GGB1532_SGB2115, Bacteroides ovatus) and Bacillota (GGB31790_SGB47293) were enriched in individuals with worse physical condition, suggesting that these taxa may serve as potential biomarkers for predicting maternal health status. Beyond taxonomic variation, we observed substantial differences in microbial functional pathways between individuals with better and worse postpartum physical condition. Individuals in better condition showed enrichment of pathways related to ornithine degradation and the TCA cycle, whereas pathways associated with glutamate degradation and sucrose degradation were depleted. This pattern suggests that individuals with better postpartum physical condition may benefit from increased availability of glutamate and its derivatives, such as GABA, to support metabolic and nutritional demands. In addition, the depletion of several carbohydrate metabolism pathways in individuals with better condition may reduce competition for nutrients between the female and gut microbiota, favoring female energy absorption. Alternatively, this depletion may stem from higher female digestive efficiency, where increased carbohydrate absorption leaves fewer resources for the gut microbiota. As noted above, further research is needed to clarify the mechanistic links between pregnancy-associated changes in gut microbiota composition, metabolic function, and physical condition.

Finally, although a more diverse gut bacterial community is often associated with better health outcomes [84, 85], contrary to our expectations, individuals with poorer postpartum physical condition exhibited higher alpha diversity and more complex microbial network structures compared with individuals in better physical condition. This suggests that greater microbial diversity may not always be beneficial during pregnancy. One possible explanation is that a more diverse and highly connected microbial community may be less functionally specialized toward supporting maternal metabolic processes or immune function during this critical period. Additionally, an increase in immune tolerance during pregnancy may benefit pathogenic bacteria that are normally excluded from the gut, both increasing gut microbial diversity while simultaneously decreasing maternal physical condition. Further studies investigating how gut microbial diversity affects host metabolic processes and health outcomes would be valuable.

## Conclusions

In conclusion, our study demonstrates that the gut microbiome of female Phayre’s leaf monkeys undergoes dynamic taxonomic and functional transitions across reproductive states, patterns that are closely associated with fluctuations in maternal hormone levels. By utilizing shotgun metagenomic sequencing, we characterized shifts in specific metabolic pathways, including those involved in amino acid and carbohydrate metabolism, that are critical during pregnancy but remain difficult to resolve using 16S amplicon sequencing approaches. Our findings link microbial dissimilarity across the reproductive cycle to health- and fitness-relevant outcomes, specifically postpartum physical condition. These results suggest that a more stable microbial profile may be associated with better postpartum health. Future research should focus on identifying the specific taxa that shift across reproductive states and investigating how microbial metabolites directly influence maternal physical condition to better understand the underlying mechanisms. Collectively, our study expands our current understanding of the interplay between the gut microbiota, maternal hormones, and maternal strategies during gestation, highlighting the potential role of the microbiome as a mediator of maternal–fetal conflict.

## List of abbreviations

*fE*: fecal estrogen
*fP*: fecal progesterone
*GDM*: gestational diabetes mellitus
*IBI*: Interbirth interval

## Declarations

### Ethics

All methods and protocols were approved by the Stony Brook University IACUC (IDs: 20041120-20051120) and complied with the applicable laws of Thailand and the United States. Permission to conduct research in Thailand was granted by the National Research Council of Thailand, the Department of National Parks, Wildlife, and Plant Conservation, and Phu Khieo Wildlife Sanctuary.

### Availability of data and material

Raw DNA sequence, rainfall, reproductive state, and hormone data for the analysis can be accessed through NCBI under BioProject ID PRJNA1453589. Code for analysis can be found at GitHub repository: https://github.com/Mallott-Lab/Leaf-Monkey-Metagenomics. Electronic supplementary material is available online at GitHub repository: https://github.com/Mallott-Lab/Leaf-Monkey-Metagenomics

### Competing interests

The authors declare no competing interests.

### Funding

Funding for this study was provided by the National Science Foundation (BCS-0542035 to AK and CB; BCS-0452635 to CB and AL), the Leakey Foundation (to AL), the American Society of Primatologists (to AL), and the National Institute of General Medical Sciences of the National Institutes of Health under award number R35GM157055 (to EKM). The content is solely the responsibility of the authors and does not necessarily represent the official views of the National Institutes of Health.

### Authors’ contributions

AL and EKM conceptualized the study. Funding was acquired by AK, CB, AL, and EKM. AK and CB provided the rainfall, body condition, and interbirth interval data. AL performed the majority of field data collection and conducted the hormone analyses. KG performed the molecular lab work. TS-N carried out the formal analysis and created the visualizations. TS-N and EKM wrote the manuscript, which was edited by AL, CB, AK, and KG. All authors gave final approval for publication and agreed to be held accountable for the work performed therein.

## Acknowledgements

The authors thank the National Research Council of Thailand, the Department of National Parks, Wildlife, and Plant Conservation, and Phu Khieo Wildlife Sanctuary for permission to conduct research in Thailand. We thank the staff of Phu Khieo Wildlife Sanctuary, including Kitti Kreetiyutanont, Mongkol Kumsuk, Kanjana Nitaya, and Jarupol Prabnasuk. We also thank Naris Bhumpakphan, Wichan Eiadthong (Kasetsart University), Warren Brokelman (Mahidol University), and Tommaso Savini (King Mongut’s University of Technology, Thonburi) for collaboration and support in Thailand. We thank our research assistants, sanctuary rangers, and Eileen Larney for help with data collection. We thank Jacinta Beehner (University of Michigan) and Nancy Czekala (formerly at the Institute for Conservation Research at the San Diego Zoo) for laboratory space to analyze hormone samples. We thank the Alvin J. Siteman Cancer at Washington University School of Medicine, Barnes-Jewish Hospital, and the Institute for Clinical and Translational Sciences (ICTS) at Washington University in St. Louis for the use of the Genome Technology Access Center (GTAC), which provided sequencing services. The authors acknowledge the Research Infrastructure Services (RIS) group at Washington University in St. Louis for providing computational resources and services.

## Supplementary data

**Figure S1.**
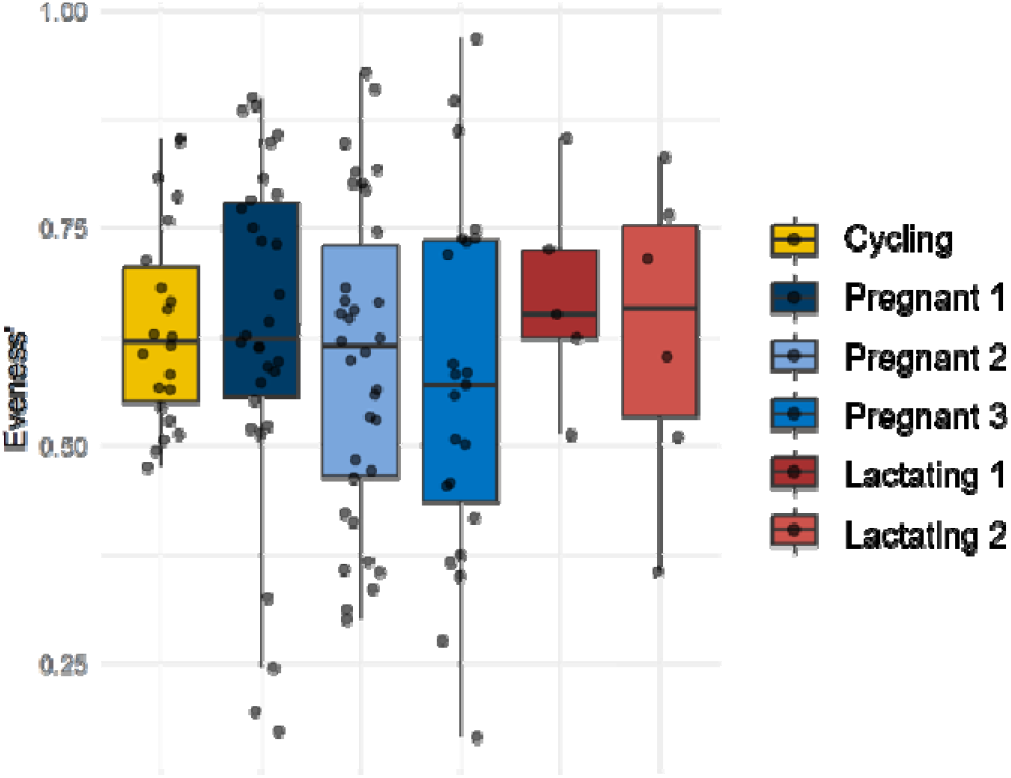
Box plots showing species evenness of gut microbiota across reproductive states. In each box plot, the center line represents the median, the box indicates the interquartile range (IQR), and the whiskers extend to 1.5 × IQR. Statistical analysis was performed using LMMs with individual identity a a random effect.

**Figure S2.**
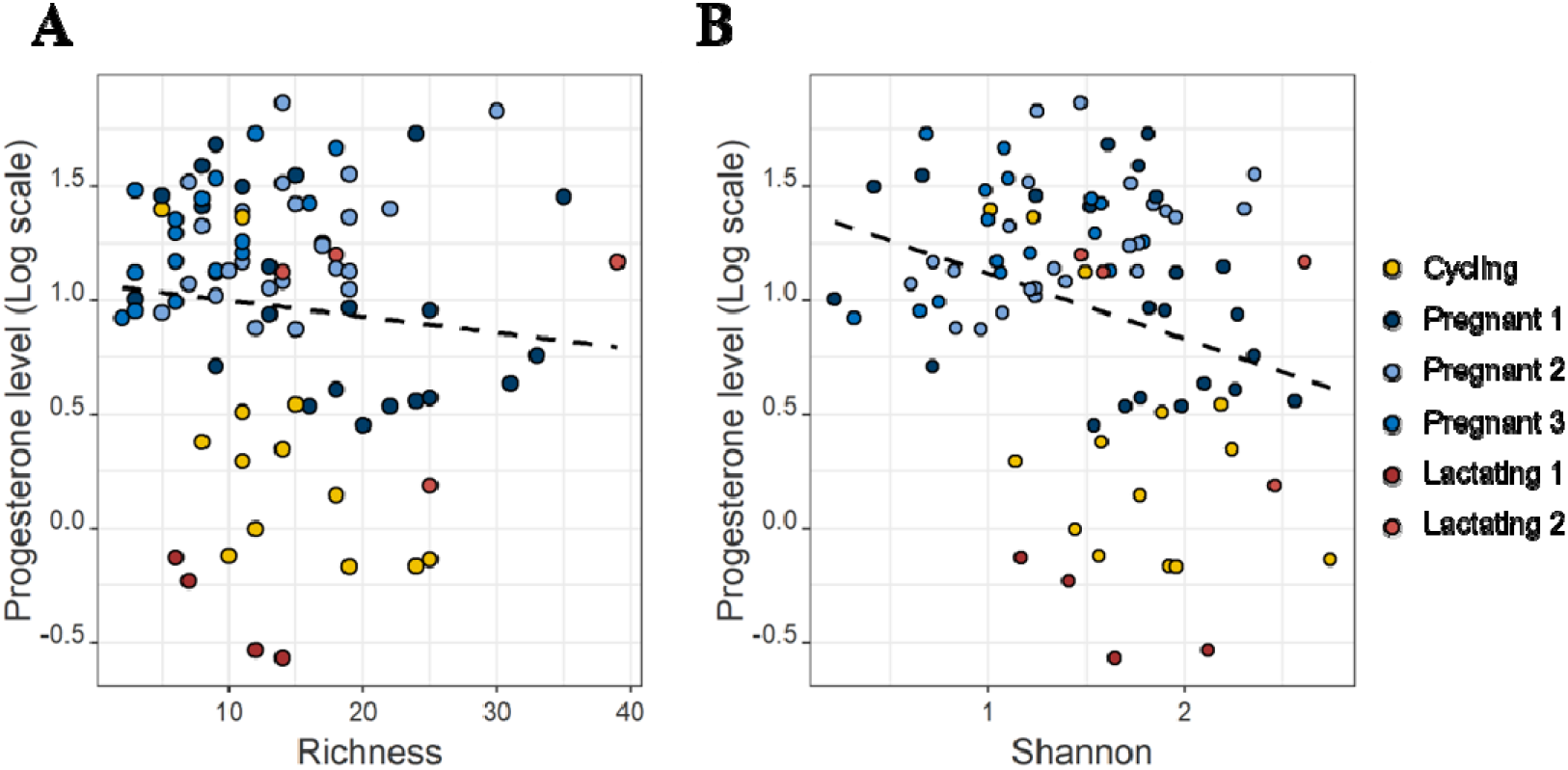
Scatter plots showing correlations between progesterone levels and (A) species richness and (B) Shannon diversity. Dotted lines represent the linear regression fit. Repeated measures correlation coefficients (*r_rm_*) were *r_rm_* = −0.256 (*p* = 0.026) for species richness, and *r_rm_* = −0.350 (*p* = 0.002) for Shannon diversity.

**Figure S3.**
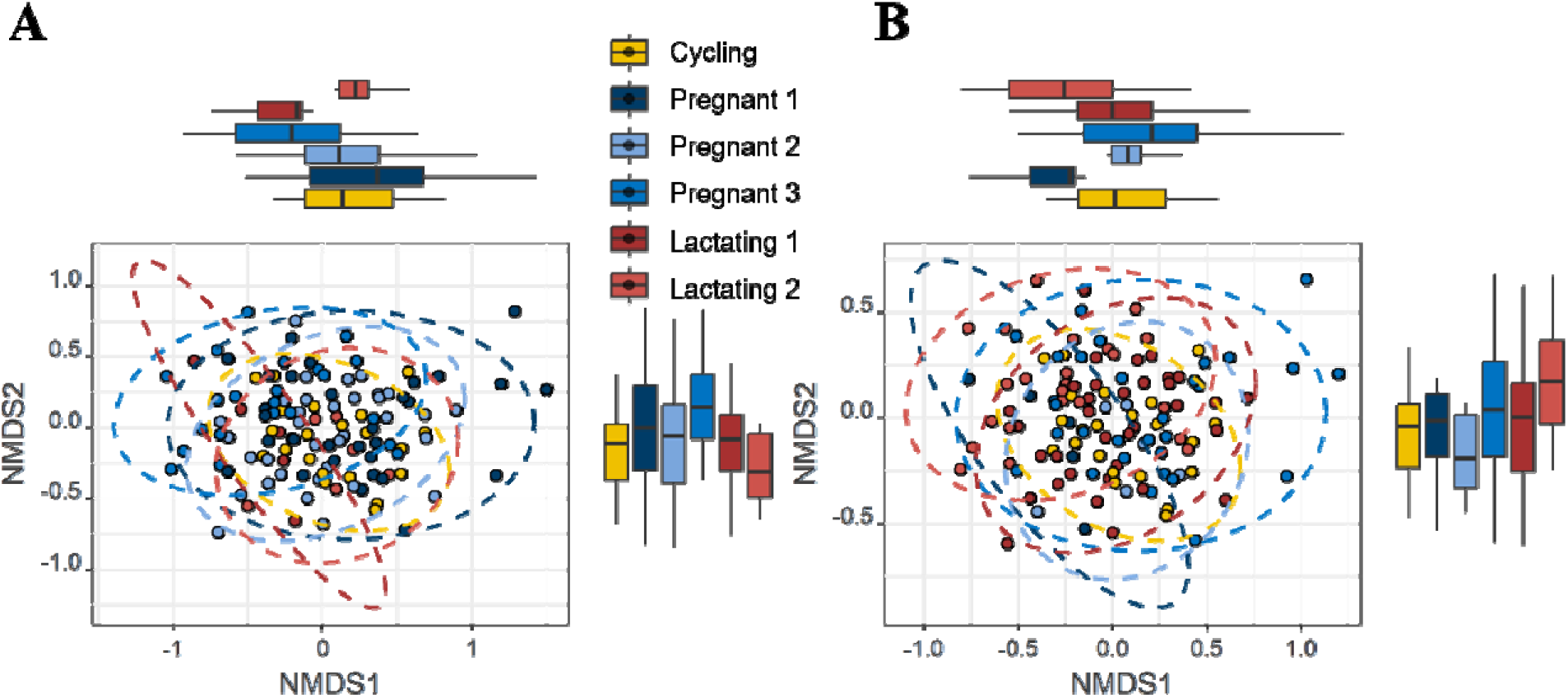
Nonlinear multidimensional scaling (NMDS) plots across reproductive states. NMDS plot based on (**A**) Bray-Curtis and (**B**) Jaccard distances of gut microbiota across reproductive states. Box plots along the top and right sides represent the distribution of NMDS1 and NMDS2 values for each reproductive state. In each box plot, the center line represents the median, the box indicates the interquartile range (IQR), and the whiskers extend to 1.5 × IQR. For the Bray-Curtis NMDS plot, stress = 0.241, *R²* for reproductive states = 0.053 (*p* = 0.010). For the Jaccard NMDS plot, stress = 0.240, *R²* for reproductive states = 0.053 (*p* = 0.027).

**Figure S4.**
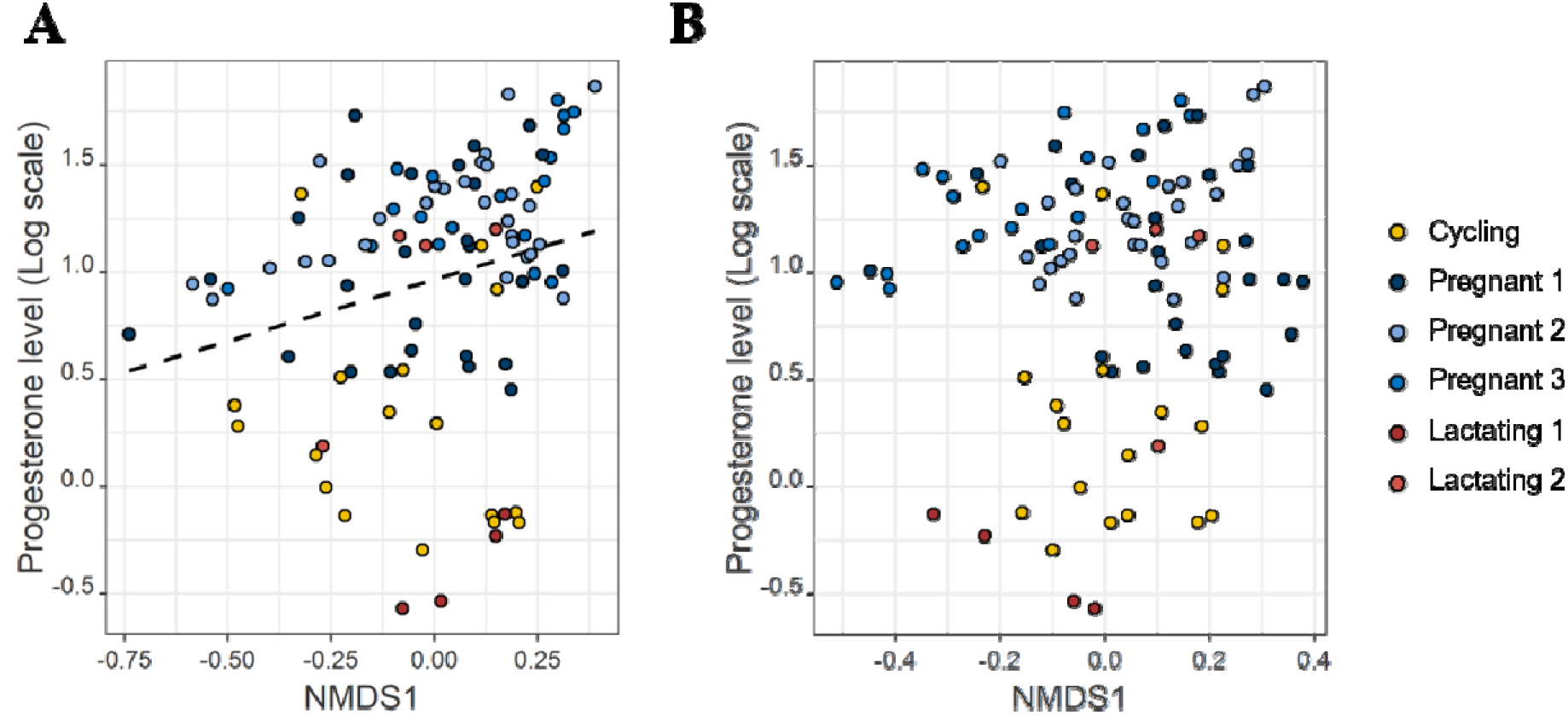
Scatter plots showing correlations between progesterone levels and NMDS1 values derived from (**A**) weighted UniFrac and (**B**) unweighted UniFrac. Dotted lines represent the linear regression fit. Repeated measures correlation coefficient (*r_rm_*) for weighted UniFrac was *r_rm_* = 0.251 (*p* = 0.017); no significant correlation was observed for unweighted UniFrac.

**Figure S5.**
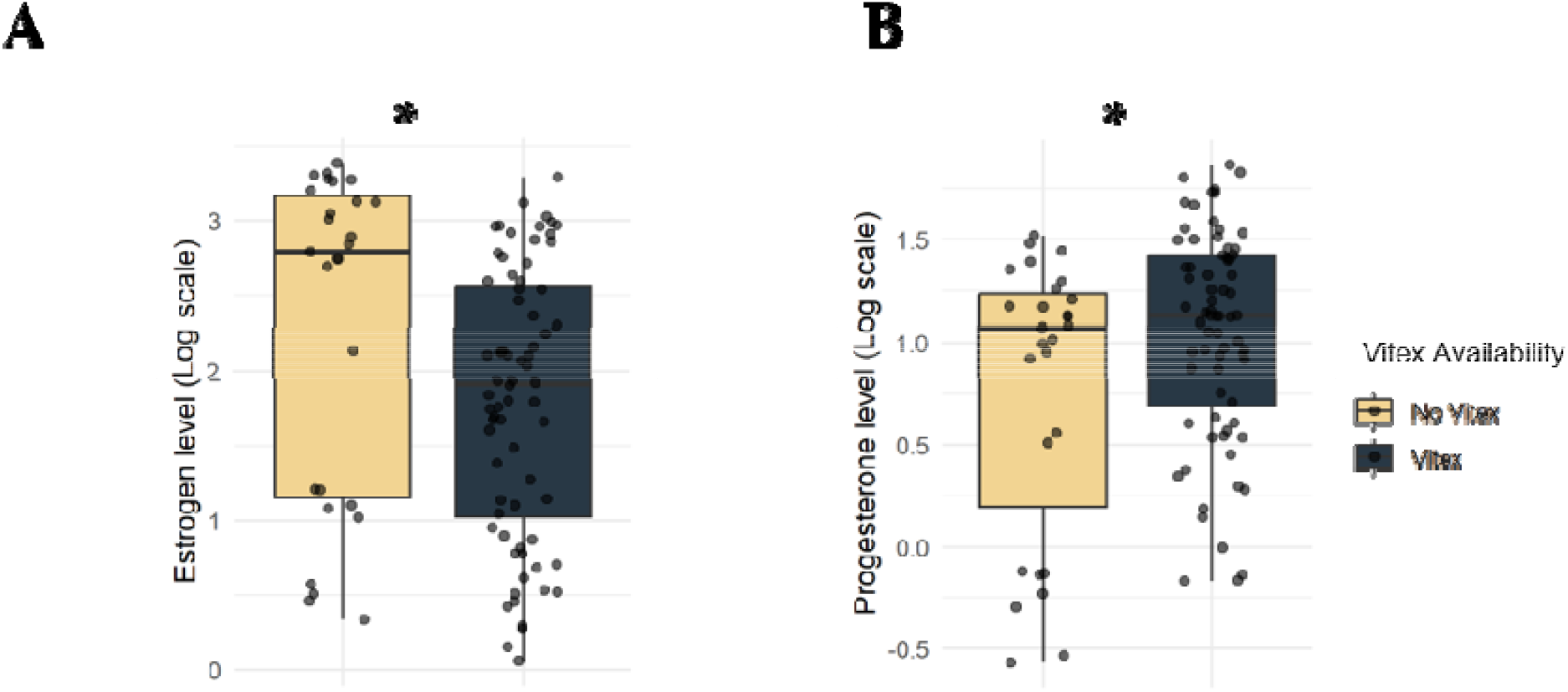
Box plots of (**A**) fE and (**B**) fP level of samples collected in the *Vitex* or non-*Vitex* season. In each box plot, the center line represents the median, the box indicates the interquartile range (IQR), and the whiskers extend to 1.5 × IQR. Statistical analysis was performed using LMMs with individual identity as a random effect. Statistical significance levels are indicated as follows: \*\**p* < 0.01.

**Figure S6.**
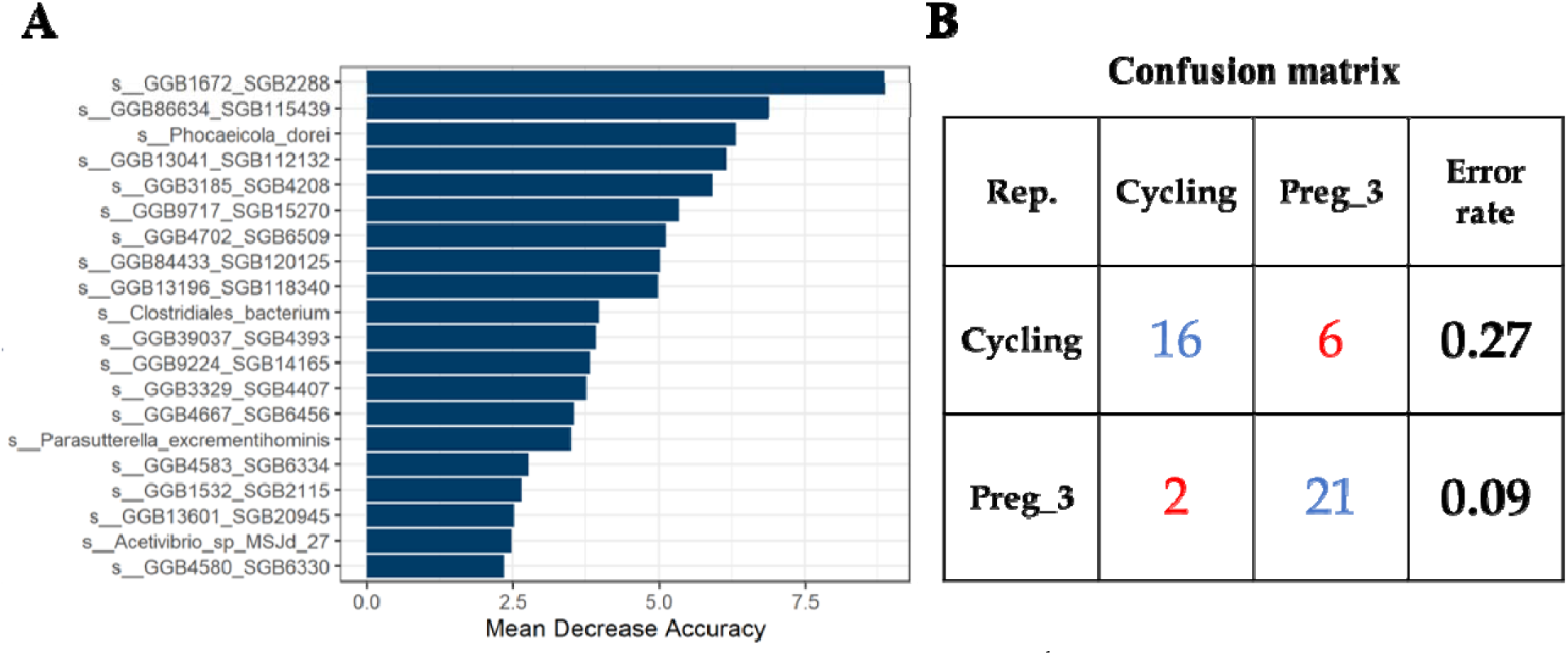
Identification of key taxa distinguishing cycling and 3^rd^ trimester using a Random Forest algorithm. (A) Bar plot showing the taxa most important for distinguishing cycling and 3^rd^ trimester, ordered by mean decrease in accuracy. (B) Confusion matrix demonstrating classification performance between cycling and 3^rd^ trimester using the random forest model.

**Figure S7.**
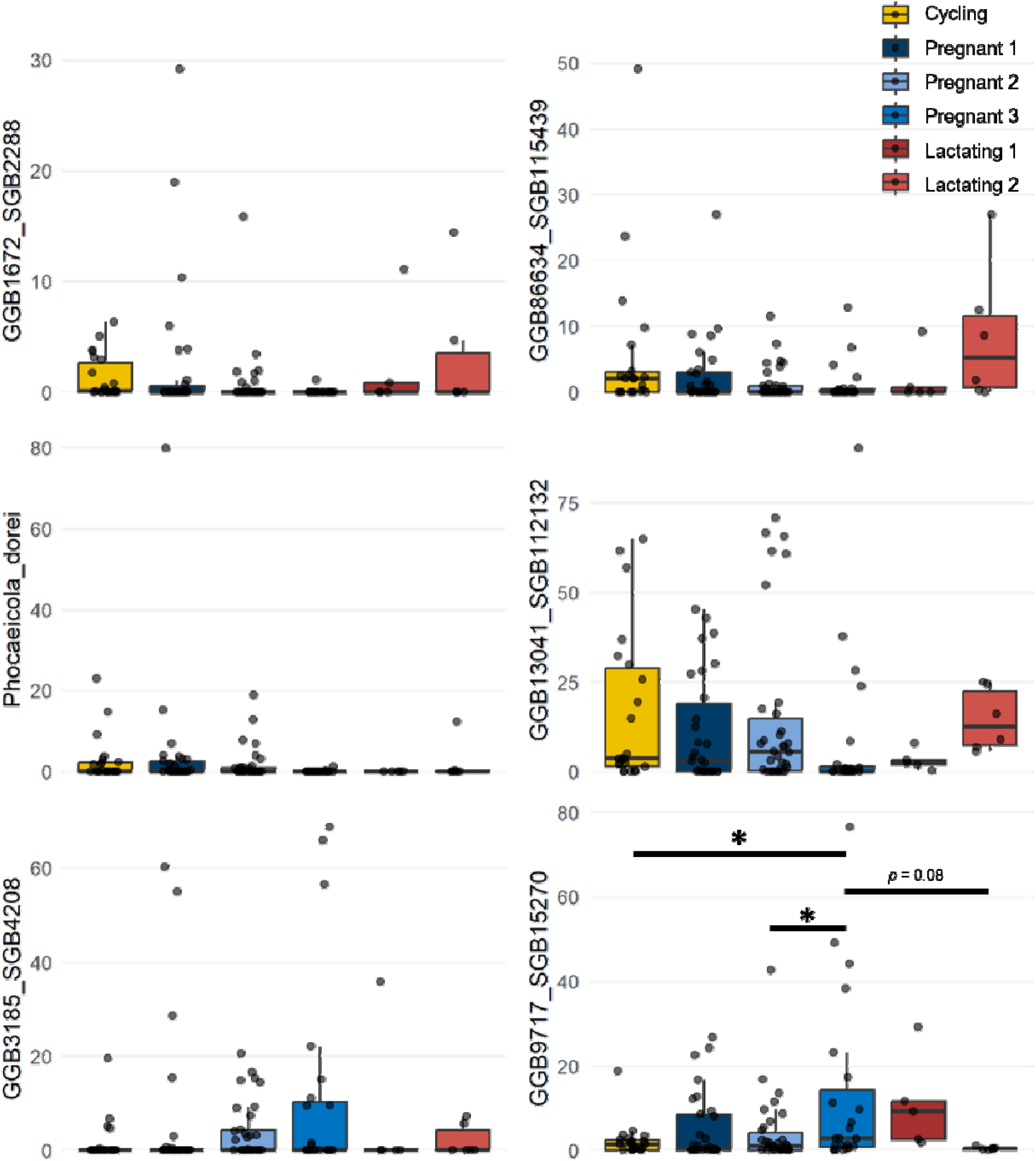
Box plots showing the relative abundance of selected taxa via Random Forest importance scores. In each box plot, the center line represents the median, the box indicates the interquartile range (IQR), and the whiskers extend to 1.5 × IQR. Statistical analysis was performed using LMMs with individual identity as a random effect, followed by Tukey’s HSD. Statistical significance level: \**p* < 0.05.

**Figure S8.**
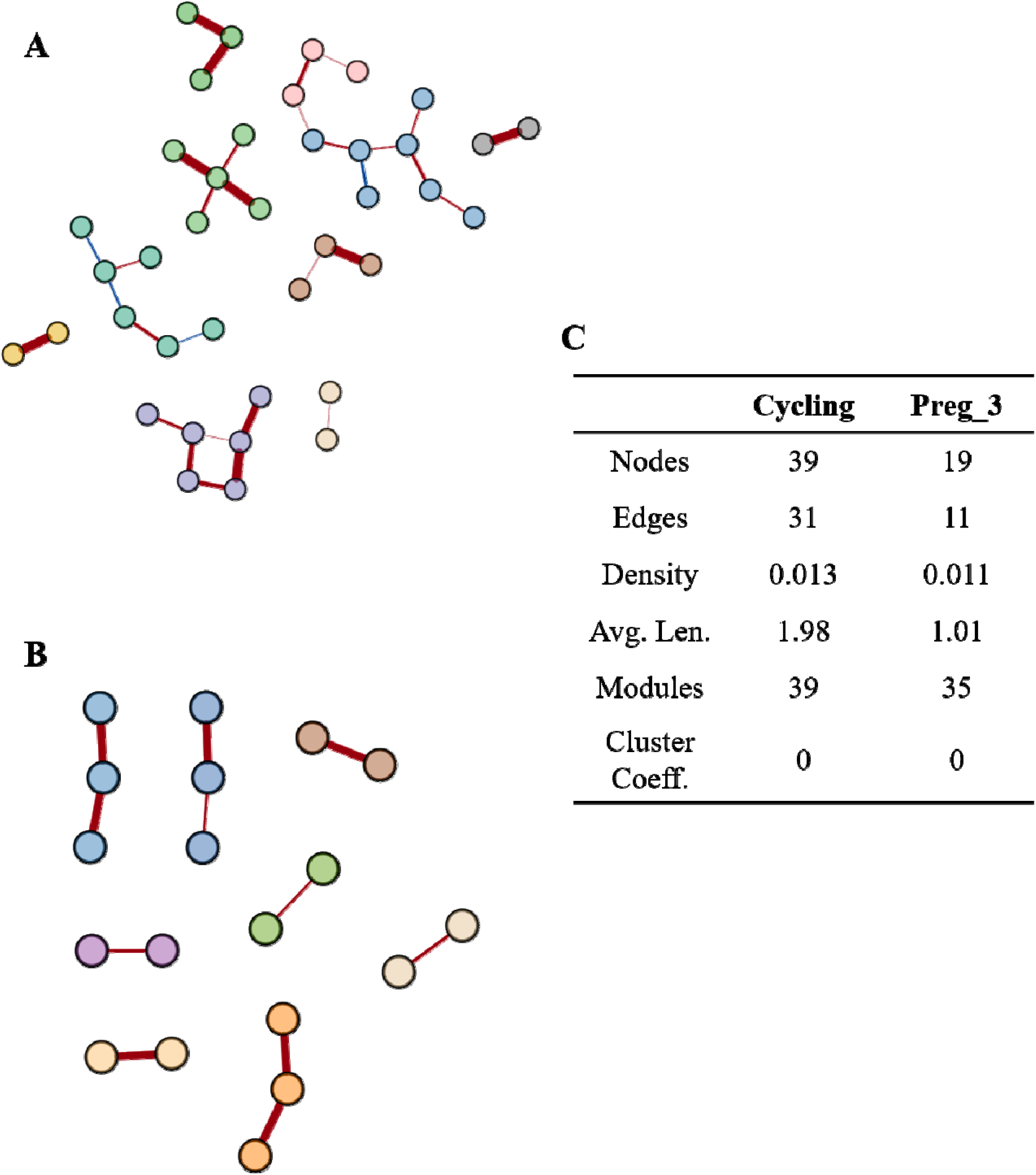
Network analysis of gut microbiota composition in (**A**) cycling and (**B**) 3^rd^ trimester. Each node represents a taxon, and each edge represents a weighted correlation. Edge thickness indicates th strength of the correlation, with red edges representing positive correlations and blue edges representing negative correlations. (C) Summary of network properties. Nodes indicate the number of taxa (nodes) in each network; Edges indicate the number of connections between taxa; Density represents the ratio of observed connections to the total possible connections; Average path length (avg. len.) indicates the mean of the shortest path lengths between all node pairs; Modules represent groups of taxa that are more densely connected to each other than to nodes outside the module; and the Clustering coefficient indicates the tendency of nodes in the network to form tightly connected groups.

**Figure S9.**
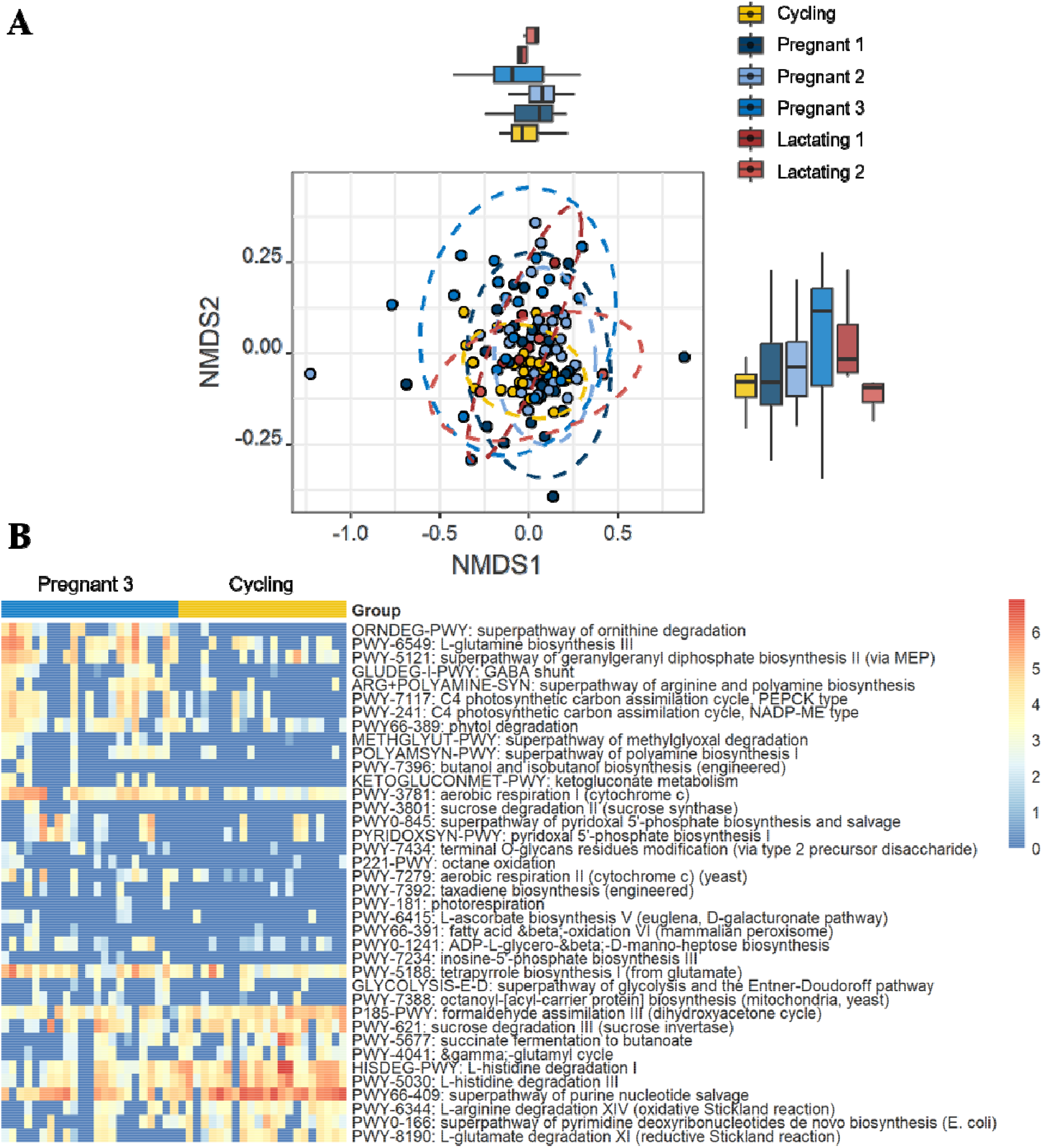
(**A**) Nonlinear multidimensional scaling (NMDS) plot based on Jaccard distances of gut microbiota pathway abundances across reproductive states. Box plots along the top and right sides show the distribution of NMDS1 and NMDS2 values for each reproductive state. Stress = 0.126; *R²* for reproductive states = 0.097 (*p* = 0.003). **(B)** Heatmap of pathways enriched or depleted in the 3^rd^ trimester compared with cycling (corresponding to Figure 3C). The color scale represents the logarithm of pathway read counts, with red indicating enrichment and blue indicating depletion in the 3^rd^ trimester. Full pathway names are shown on the right side of the plot.

**Figure S10.**
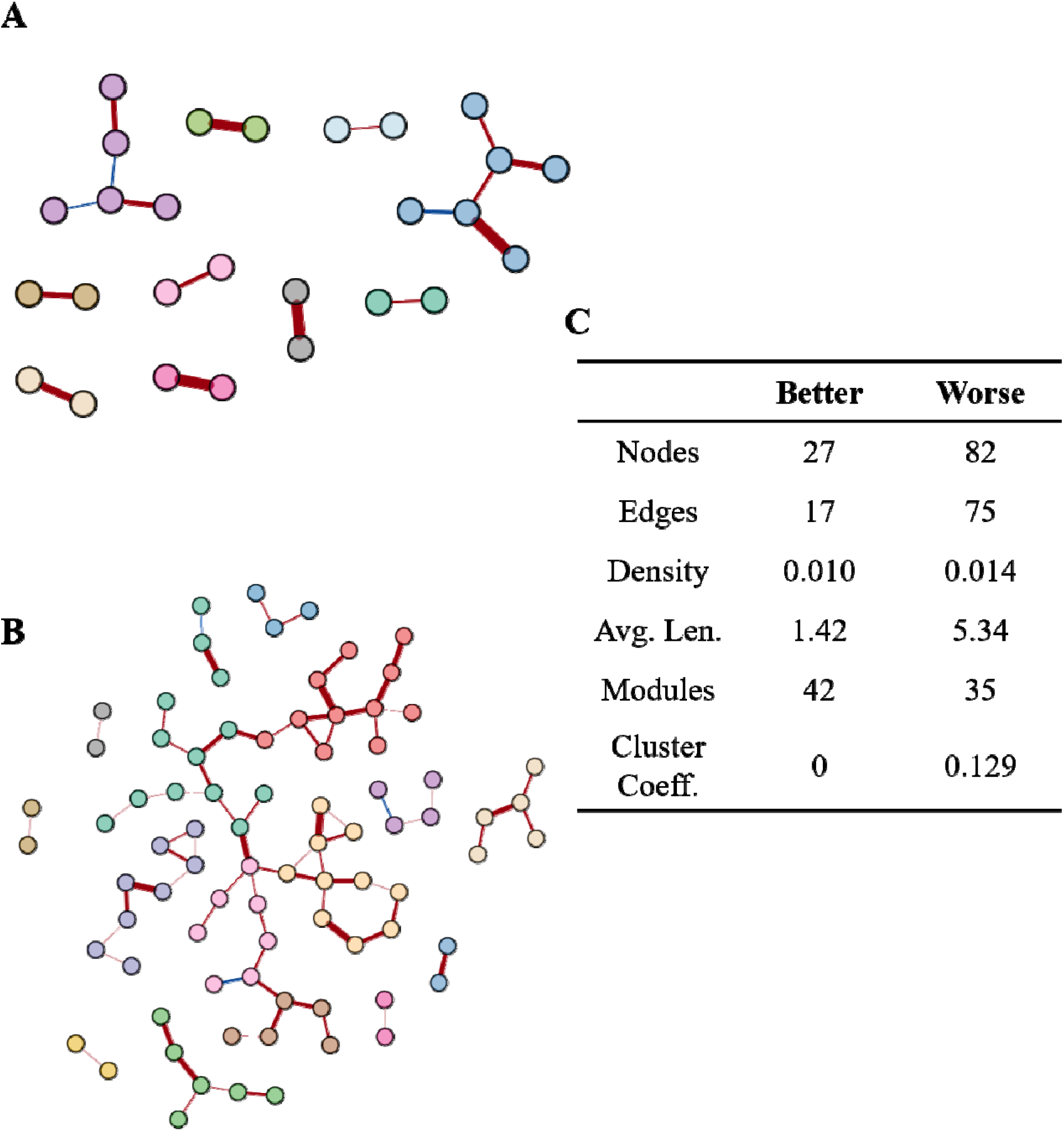
Network analysis of gut microbiota composition in individuals with (**A**) better and (**B**) worse postpartum physical condition. Each node represents a taxon, and each edge represents a weighted correlation. Edge thickness indicates the strength of the correlation, with red edges representing positive correlations and blue edges representing negative correlations. (**C**) Summary of network properties. Nodes indicate the number of taxa (nodes) in each network; Edges indicate the number of connection between taxa; Density represents the ratio of observed connections to the total possible connections; Average path length (avg. len.) indicates the mean of the shortest path lengths between all node pairs; Modules represent groups of taxa that are more densely connected to each other than to nodes outside the module; and the Clustering coefficient indicates the tendency of nodes in the network to form tightly connected groups.

**Figure S11.**
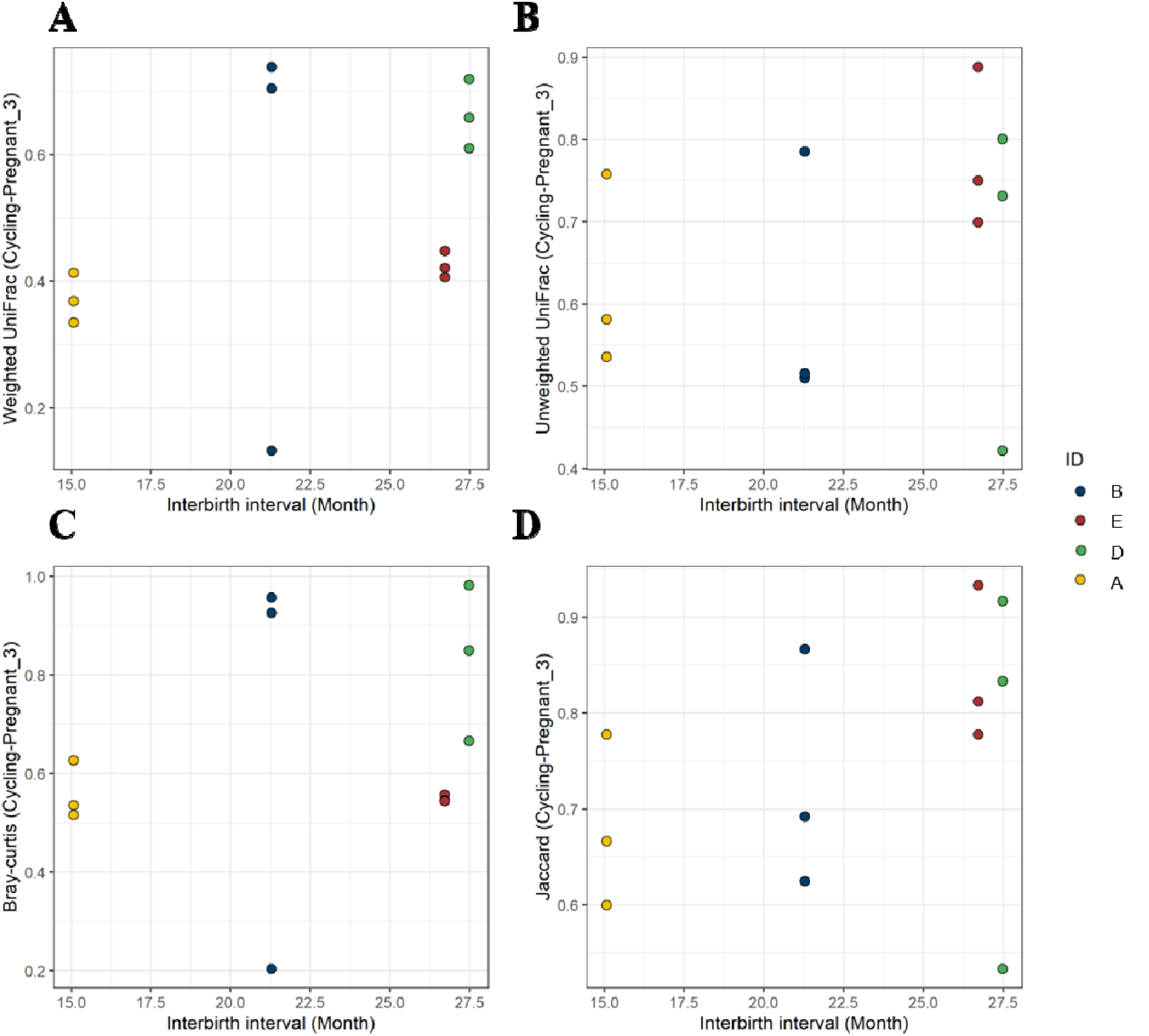
Scatter plots showing correlations between interbirth interval (IBI) and species dissimilarity between the cycling and 3^rd^ trimester. Species dissimilarity was calculated using (**A**) weighted UniFrac ( = 0.39, *p* = 0.205), (**B**) unweighted UniFrac (*r* = 0.28, *p* = 0.373), (**C**) Bray–Curtis (*r* = 0.25, *p* = 0.434), and (**D**) Jaccard distances (*r* = 0.40, *p* = 0.202). Each ID represents an individual female.

**Figure S12.**
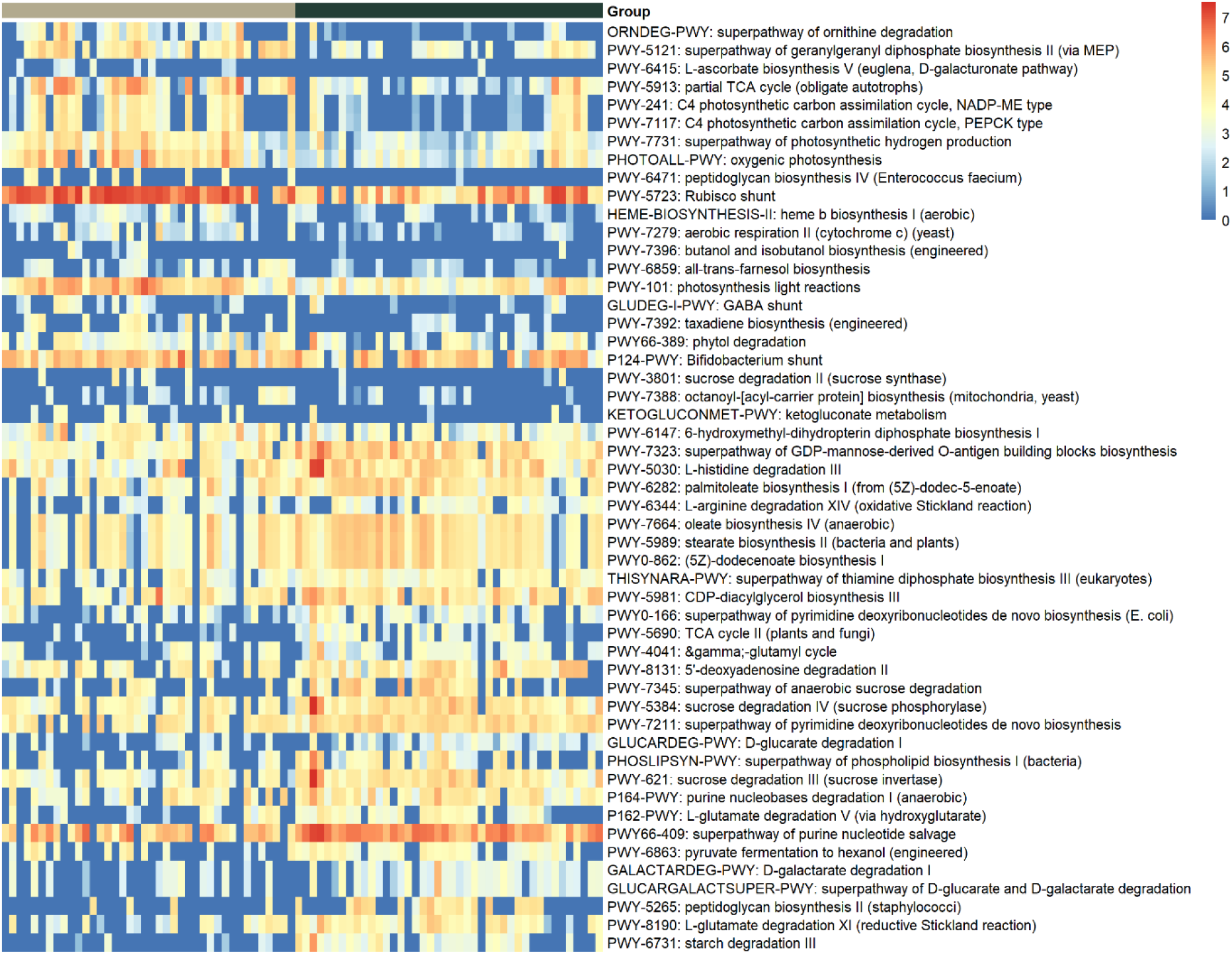
Heatmap of pathways enriched or depleted in individuals with better compared to worse physical condition (corresponding to Figure 5B). The color scale represents the logarithm of pathway read counts, with red indicating enrichment and blue indicating depletion in the better physical condition. Full pathway names are shown on the right side of the plot.

## Notes

https://github.com/Mallott-Lab/Leaf-Monkey-Metagenomics

### Competing Interest Statement

The authors have declared no competing interest.

